# A hierarchical transcriptional network activates specific CDK inhibitors that regulate G2 to control cell size and number in Arabidopsis

**DOI:** 10.1101/2021.10.16.464643

**Authors:** Yuji Nomoto, Hirotomo Takatsuka, Kesuke Yamada, Toshiya Suzuki, Takamasa Suzuki, Ying Huang, David Latrasse, Jing An, Magdolna Gombos, Christian Breuer, Takashi Ishida, Kenichiro Maeo, Miyu Imamura, Takafumi Yamashino, Keiko Sugimoto, Zoltán Magyar, László Bögre, Cécile Raynaud, Moussa Benhamed, Masaki Ito

**Affiliations:** School of Biological Science and Technology, College of Science and Engineering, Kanazawa University, Kakuma-machi, Kanazawa, 920-1192 Japan; National Institute of Genetics, 1111 Yata, Mishima, Shizuoka 411-8540, Japan; College of Bioscience and Biotechnology, Chubu University, Kasugai, Aichi 487-8501, Japan; Université Paris-Saclay, CNRS, INRAE, Univ Evry, Institute of Plant Sciences Paris-Saclay (IPS2), 91405, Orsay, France; Institute of Plant Biology, Biological Research Centre, Szeged, 6726, Hungary; RIKEN Center for Sustainable Resource Science, Yokohama 230-0045, Japan; Graduate School of Bioagricultural Sciences, Nagoya University, Furocho, Chikusa-ku, Nagoya, 464-8601, Japan; Department of Biological Sciences, Graduate School of Science, The University of Tokyo, 7-3-1 Hongo, Bunkyo-ku, Tokyo 113-0033, Japan; Royal Holloway University of London, Department of Biological Sciences, Centre for Systems and Synthetic Biology, Egham, TW20 0EX, UK

## Abstract

How cell size and number are determined during organ development remains a fundamental question in cell biology. Here, we identified a GRAS family transcription factor, called SCARECROW-LIKE28 (SCL28), with a critical role in determining cell size in Arabidopsis. SCL28 is part of a transcriptional regulatory network downstream of the central MYB3Rs that regulate G2 to M phase cell cycle transition. We show that SCL28 forms a dimer with the AP2-type transcription factor, AtSMOS1, which defines the specificity for promoter binding and directly activates transcription of a specific set of SIAMESE-RELATED (SMR) family genes, encoding plant-specific inhibitors of cyclin-dependent kinases and thus inhibiting cell cycle progression at G2 and promoting the onset of endoreplication. Through this dose-dependent regulation of *SMR* transcription, SCL28 quantitatively sets the balance between cell size and number without dramatically changing final organ size. We propose that this hierarchical transcriptional network constitutes a cell cycle regulatory mechanism that allows to adjust cell size and number to attain robust organ growth.

---

During organ growth and development, cell proliferation is intricately controlled in space and time. Governed by developmental programs (Roeder et al., 2010; Polyn et al., 2015; Andriankaja et al., 2012) and influenced by active plant responses to environmental conditions (Komaki and Sugimoto, 2012; Qi and Zhang, 2020), the number of cells produced during organ growth is set by the cell cycle speed during proliferation and the point when cells exit to cellular differentiation. Controlled cell proliferation requires coordinately regulated gene expression within and upon exit from the cell cycle. Generally, there are two main groups of genes showing waves of transcription during the cell cycle; G1/S-specific and G2/M-specific genes (Berckmans and De Veylder, 2009). The G1/S-specific genes facilitate initiation and progression of DNA replication and are typically regulated by the activity of E2F transcription factors (Vandepoele et al., 2005; Őszi et al., 2020). Generally, E2F dimerizes with Dimerization Partner (DP) proteins to activate or repress their target genes, depending on their association with the Retinoblastoma-Related (RBR) repressor protein (Desvoyes and Gutierrez, 2020). On the other hand, most G2/M-specific genes are positively or negatively regulated by MYB3R family of transcription factors in plants (Haga et al., 2011; Kobayashi et al., 2015). Some members of MYB3Rs are specifically expressed during G2/M and act as transcriptional activators, while others act as transcriptional repressors of G2/M-specific genes (Ito et al., 2001; Haga et al., 2007; Okumura et al., 2021). Recent studies identified the two main groups of transcription factors, E2Fs and MYB3Rs, which had been studied separately, as part of the same multi-protein complex in Arabidopsis (Kobayashi et al., 2015). This E2F-MYB3R complex is evolutionarily related to the DREAM (DP, Retinoblastoma-like, E2F, and MuvB) complex reported in *Drosophila* and human cells (Kobayashi et al., 2015; Ning et al., 2020; Lang et al., 2021). The metazoan DREAM complex plays a predominant role in repressing both G1/S- and G2/M-specific genes, thus promoting cell cycle exit and maintaining cellular quiescence (Sadasivam and DeCaprio, 2013; Fischer and Müller, 2017). The DREAM complex in Arabidopsis shows significant differences from metazoan complexes, which include involvement of plant-specific subunits and existence of diversified complexes with different subunit compositions (Kobayashi et al., 2015; Ning et al., 2020; Desvoyes and Gutierrez, 2020).

Transcriptional regulation during cell cycle generally constitutes multi-layered hierarchical networks, in which master regulators regulate other transcription factors, which further regulate each other or downstream genes (Lee et al., 2002; Zhu et al., 2004; Millour et al., 2011). Notably, studies in yeasts showed that cell cycle transcriptional activators that function during one stage of the cell cycle regulate transcriptional activators that function during the next stage, forming a connected regulatory network that is itself a cycle (Simon et al., 2001). However, in plants, such a hierarchical network composed of cell cycle transcription factors, E2F and MYB3R, has not yet been uncovered. Exploring the transcription network during cell cycle may uncover missing important factors and hidden mechanisms governing plant-specific cell cycle regulation. In accordance with our work, recently a mitosis specific GRAS family transcription factor, called SCARECROW-LIKE28 (SCL28), has been identified as being directly regulated by MYB3Rs (Goldy et al., 2021). Here we demonstrate that SCL28 acts in association with the AP2-type transcription factor, AtSMOS1, to directly activate transcription of a specific set of *SMR* family genes, encoding plant-specific cyclin-dependent kinase (CDK) inhibitors (Kumar et al., 2015). This regulatory network inhibits the G2 to M phase transition during the cell cycle and promotes the onset of endoreplication, an atypical cell cycle consisting of repeated DNA replication without mitosis (Komaki and Sugimoto, 2012). Our study identified a G2/M regulatory pathway that controls cell cycle length, and is likely to be important in optimizing cellular functions by setting cell size and number during organ growth.

## Results

### Identification of a mitotic GRAS-type transcription factor

By analyzing transcripts specific in mitotic cells, we have sience identified an Arabidopsis GRAS family transcription factor that we designated *E1M* (Ito et al., 1999). A recent report looking for mitotic genes in the root meristem uncovered the same gene called *SCL28* (Goldy et al., 2021), the name hereafter also adopted in this study. In a synchronized culture of Arabidopsis MM2d cells, we showed that this gene exhibited G2/M-specific transcript accumulation, which closely resembles that of the mitotic cyclin *CYCB1*;*2* (Fig. 1a). As for most G2/M-specific genes, the so-called mitosis-specific activator (MSA) element that serves as a binding motif for MYB3Rs (Ito et al., 1998; Haga et al., 2011) were repeatedly present within the proximal promoter regions of *SCL28* (Fig. 1b). Binding of MYB3R to the *SCL28* promoter is supported by our published data from chromatin immunoprecipitation (ChIP) with MYB3R3 followed by high-throughput sequencing (ChIP-Seq; Fig. 1b) (Kobayashi et al., 2015) and DNA affinity purification sequencing (DAP-Seq) data reported for MYB3R1, MYB3R4, and MYB3R5 (Supplementary Fig. 1a) (O’Malley et al., 2016). In addition, loss of MYB3R activators (*myb3r1/4* double mutant) or MYB3R repressors (*myb3r1/3/5* triple mutant) resulted in significant down- or up-regulation of *SCL28*, respectively (Fig. 1c). GUS reporter activity driven by *SCL28* promoter decreased significantly in *myb3r1/4* double mutant and almost diminished by elimination of MSA elements (Fig. 1d). Collectively, these observations support the idea that *SCL28* is a direct target of both activator and repressor type MYB3Rs in *Arabidopsis*. Similar to the G2/M-specific CYCB1;1-GFP accumulation (Ubeda-Tomás et al., 2009), we observed patchy pattern of SCL28-GFP signal from a construct driven by native *SCL28* promoter in root meristem, suggesting cell cycle-regulated protein accumulation (Fig. 1e). Taken together, as has been recently shown (Goldy et al., 2021), SCL28 is a mitotic gene directly regulated by MYB3Rs.

**Fig. 1.**
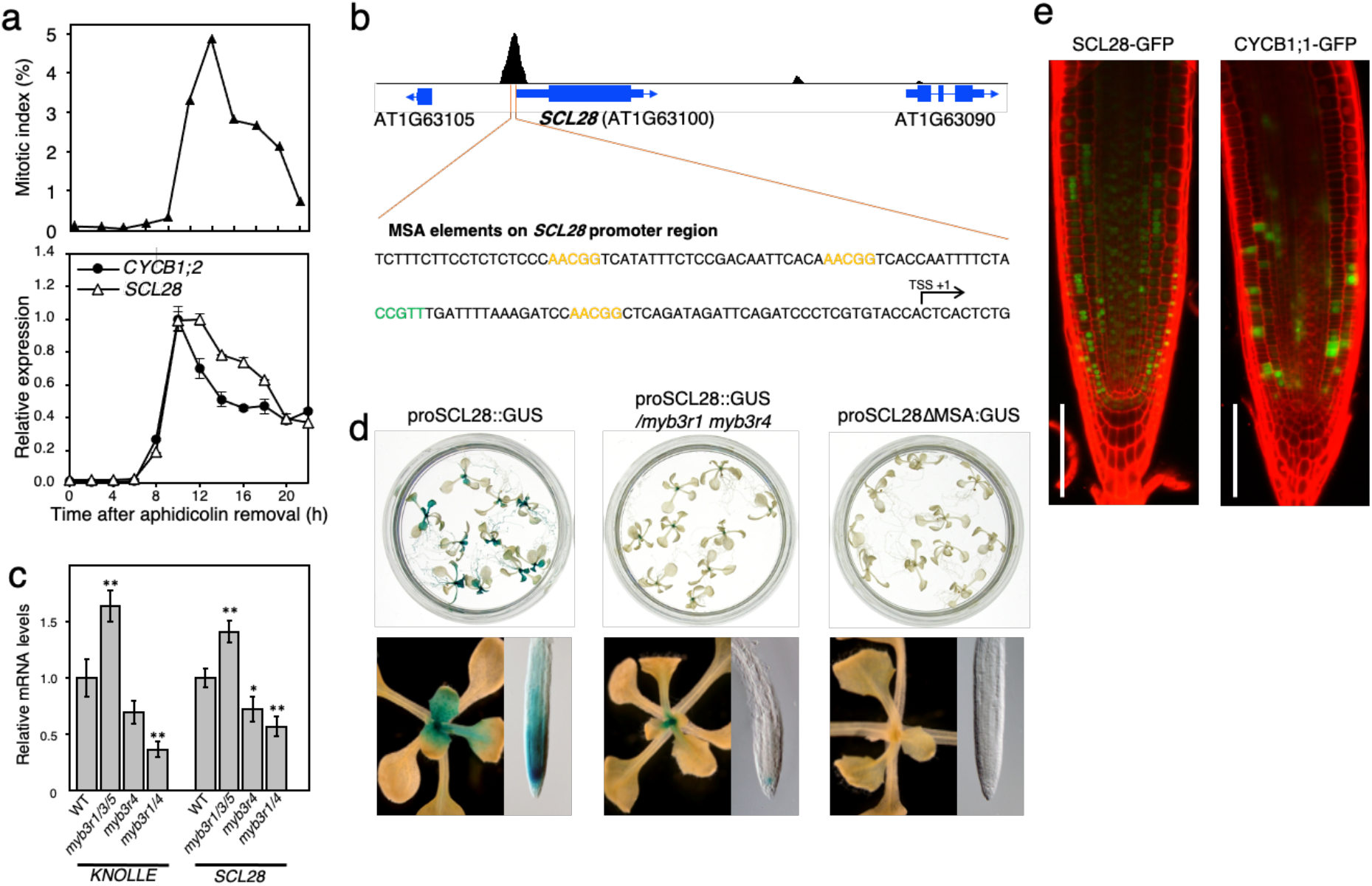
*SCL28* expression is cell cycle-regulated under the control of MYB3R transcription factors. **a** G2/M-specific accumulation of *SCL28* transcript. *Arabidopsis* MM2d cells were synchronized by aphidicolin treatment, and used for qRT-PCR analysis to quantify *SCL28* mRNA levels (lower). Synchronous progression of cell cycle was monitored by measuring mitotic index (upper). As a representative of G2/M-specific genes, *CYCB1*;*2* was also analyzed in the same way for comparison. Expression data are shown as mean ± SD (*n* = 3). **b** MYB3R3 binds to the upstream region of *SCL28*. The ChIP-Seq profile of MYB3R3 indicates its direct binding to the proximal promoter region of *SCL28*. The nucleotide sequence around the ChIP-Seq peak contains MSA elements repeated four times, of which core motives (AACGG) are shown by orange (forward orientation) and green (reverse orientation) letters. **c** Levels of *SCL28* transcripts are significantly affected by mutations in *myb3r* genes. Transcript levels of *SCL28* and *KNOLLE* were analyzed by qRT-PCR in plants 12 days after sowing (DAS) that carry mutations of *myb3r* genes in the indicated combinations (mean ± SD; *n* = 3). Statistical significance was determined using Student’s t-test. \**P* < 0.05, \*\**P* < 0.01. **d** Promoter activity of *SCL28* requires the presence of MYB3R activators and MSA elements. GUS reporter expression was analyzed in wild type (WT) and *myb3r1/4* plants carrying proSCL28::GUS and WT plants carrying proSCL28ΔMSA::GUS, in which all MSA elements in the *SCL28* promoter were mutated. GUS staining around shoot apex and root tip regions is shown at higher magnification in the lower panels. **e** Patchy pattern expression of SCL28-GFP protein. Root meristems of proSCL28::SCL28-GFP and proCYCB1;1::CYCB1;1-GFP plants were analyzed by confocal laser scanning microscopy (CLSM) after counterstaining of the cell wall with propidium iodide (PI). Green and red signals indicate fluorescence of GFP and PI, respectively. Scale bar indicates 100 *μ*m.

To evaluate whether MYB3Rs act as part of DREAM complex on *SCL28* transcription, we searched lists of target genes bound by potential DREAM components, RBR and TESMIN/TSO1-LIKE CXC 5 (TCX5), as defined by ChIP-Seq experiments reported previously (Bouyer et al., 2018; Ning et al., 2020). This examination revealed that TCX5, but not RBR, showed a significant association to *SCL28* locus *in vivo*. However, *SCL28* showed no significant change in expression in neither *rbr* nor *tcx5 tcx6* double or *e2fa e2fb e2fc* triple mutants, indicating that *SCL28* transcription depends exclusively on MYB3R (Supplementary Fig. 1b).

### *SCL28* strongly affects cell size

To analyze the biological function of SCL28, we generated transgenic plants overexpressing *SCL28* under the strong *RPS5A* promoter (proRPS5A::SCL28). These plants, herein designated *SCL28*^OE^, showed general growth retardation both in seedling and adult stages (Fig. 2a). Cell size in these plants was significantly enlarged in all examined organs and tissues, such as root tip, leaf mesophyll, embryo, and inflorescence stem (Fig. 2b and Supplementary Fig. 2). An increased cell size was apparent in both post-mitotic differentiated cells and proliferating cell populations in root and shoot apical meristems (Fig. 2c and Supplementary Fig. 2). As described later, the increased cell size in *SCL28*^OE^ leaves was associated with elevated levels of cellular ploidy induced by enhanced endoreduplication. The loss-of-function *scl28* mutant (Supplementary Fig. 3) showed an opposite cellular phenotype to *SCL28*^OE^, having cells with significantly reduced size (Fig. 2b, c and Supplementary Fig. 2). However, unlike *SCL28*^OE^, the overall stature of *scl28* mutant plants was largely normal and indistinguishable from that of wild type (WT) plants (Fig. 2a).

**Fig. 2.**
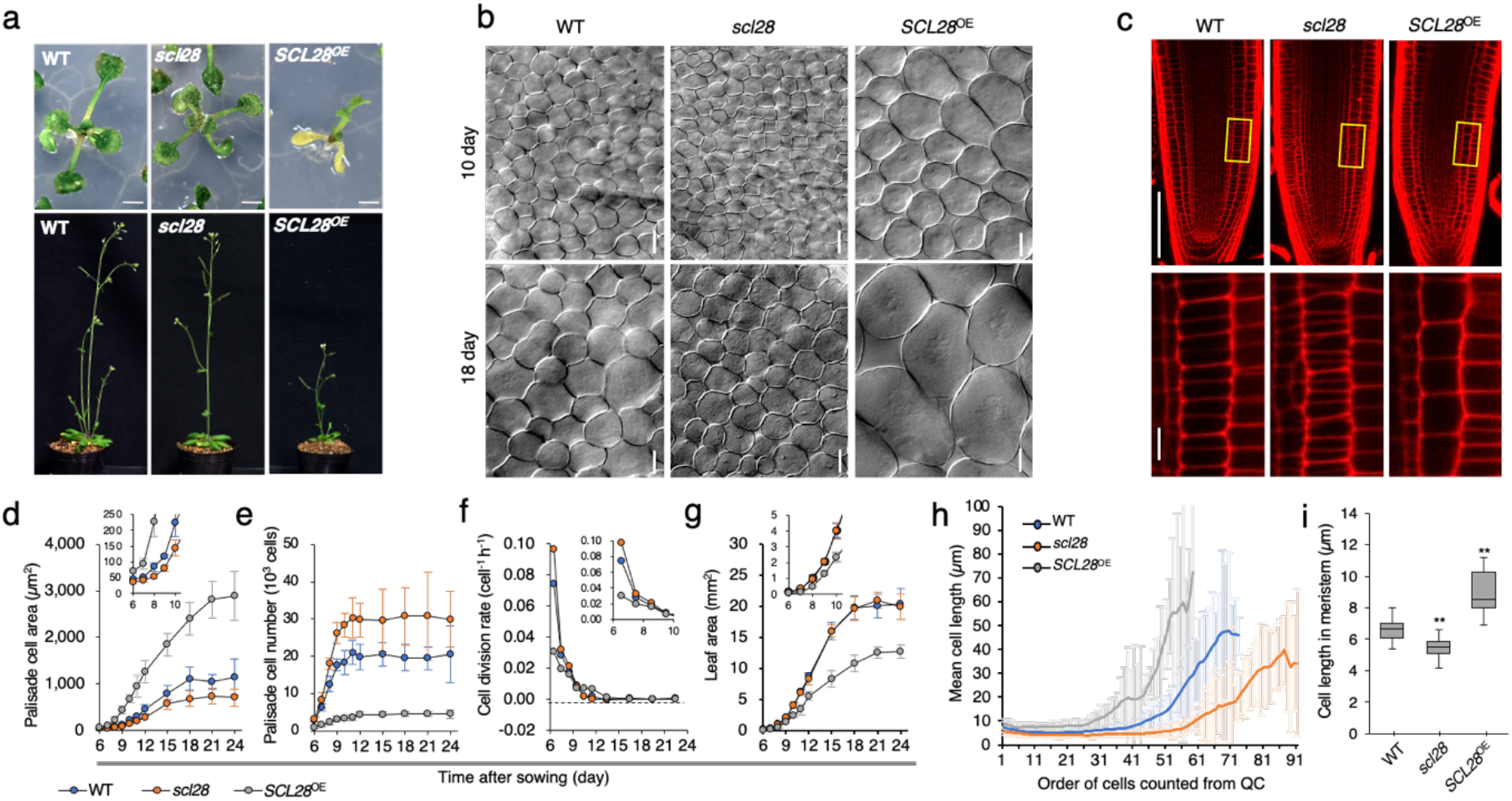
SCL28 strongly affects cell size. **a** Macroscopic phenotypes caused by loss- and gain-of-function of *SCL28*. WT, *scl28*, and *SCL28*^OE^ plants were photographed at 11 (upper) and 30 (lower) DAS. Scale bar in upper panels indicates 10 mm. **b** Cell size analysis of leaf palisade tissue from WT, *scl28*, and *SCL28*^OE^ plants. First leaf pairs from plants at 10 or 18 DAS were cleared and analyzed by differential interference contrast (DIC) microscopy. Scale bar indicates 20 *μ*m. **c** Cell size analysis of root meristem from WT, *scl28*, and *SCL28*^OE^ plants at 5 DAS. Primary roots were stained with PI, and analyzed by CLSM. Regions surrounded by yellow rectangles are shown at a higher magnification in the lower panels. Scale bars indicate 100 *μ*m (upper) and 10 *μ*m (lower). **d-g** Kinematic analysis of leaf growth in WT, *scl28*, and *SCL28*^OE^ plants. Time-course changes of palisade cell area (**d**), number of palisade cells per leaf (**e**), cell division rate (**f**), and leaf area (**g**) were analyzed in developing first leaf pairs. Data are shown as mean ± SD (*n* = 10). Inset shows the same data with an expanded *y*-axis scale. **h** Spatial distribution of cell size across root meristems in WT, *scl28*, and *SCL28*^OE^ plants at 5 DAS. Mean cell length was calculated at each position along the cortical cell file, which is defined by counts of cortical cells from the quiescent center (mean ± SD; *n* = 21). **i** Quantitative analysis of cortical cell length within root meristems. Data are shown as boxplot, where ‘boxes’ represent the interquartile range (IQR) and whiskers extend to the largest and smallest observations within 1.5 × IQR. Statistical significance compared with WT was determined using Student’s t-test. \**\*P* < 0.01 (*n* = 20).

To explore the developmental origin of cell size differences, we performed kinematic analysis on growing first leaf pairs by monitoring size and number of palisade cells (Fig. 2d–g). This analysis showed that cells in *scl28* leaves were smaller than WT already during initial stages of organ development when most cells are actively proliferating (Fig. 2d). During this early proliferative stage, increasing cell number was also more rapid in *scl28* than WT, consistent with a higher cell division rate in *scl28* leaves compared with WT (Fig. 2e, f). Conversely, duration of cell proliferation remained largely unchanged between *scl28* and WT (Fig. 2e, f). Therefore, the increased cell number in *scl28* leaves is due to accelerated cell division, rather than increased duration of cell proliferation. When leaf area was compared, however, *scl28* and WT showed no clear difference throughout the course of leaf development (Fig. 2g). In *scl28* leaves, accelerated cell division was balanced by reduced cell size, thus maintaining total organ size unchanged during leaf development. The cellular effect of *SCL28*^OE^ was generally opposite to the *scl28* mutant. The number of palisade cells per leaf was significantly reduced in *SCL28*^OE^ plants (Fig. 2e) due to severely inhibited cell proliferation (Fig. 2f). Although dramatic cell enlargement partially counteracted the reduced cell number, leaf area was still reduced in *SCL28*^OE^ compared with WT (Fig. 2d, g).

In the root tip, cell size difference was similarly apparent among *SCL28*^OE^, *scl28*, and WT (Fig. 2c, h, i). This cell size difference was maintained along the apical-basal axis of the root from cells adjacent to the quiescent center to those at the transition zone, as well as in differentiated cells at distal positions (Fig. 2h). Therefore, the behavior of cells in the root meristems of *SCL28*^OE^ and *scl28* was generally consistent with those observed in developing leaves.

### Effects of SCL28 on mitotic cell cycle and endocycle

To directly explore the role of SCL28 in cell cycle progression, we performed live-cell imaging of root meristem cells after introgression of the PCNA-GFP cell cycle marker into *SCL28*^OE^ and *scl28* mutant. The PCNA-GFP line allows quantification of cell cycle stages based on its intracellular fluorescent patterns (Yokoyama et al., 2016). During G1, PCNA-GFP appears as uniform fluorescence within the whole nuclei, which alters into dotty and speckled nuclear signals during early and late S phase, respectively. During G2, fluorescence again becomes uniform within the nuclei, then disappears upon onset of mitosis and remains undetectable until the exit from mitosis. By our definition, the G1 period begins with re-appearance of nuclear fluorescence after mitosis and ends with emergence of dotty GFP signals, whereas the G2 period begins with conversion of dotty into uniform nuclear PCNA-GFP signals and ends with disappearance of this nuclear fluorescence. Based on these definitions, we analyzed data from live-cell imaging and calculated the average duration of each cell cycle phase in WT root meristem cells to be 5.1 h in G1, 2.5 h in S, and 11.3 h in G2 (Fig. 3a, b). G2/M duration was oppositely affected in the *scl28* and *SCL28*^OE^ lines, shortened 19% in the former and lengthened 33% in the later. In contrast, G1 length showed only limited alteration, but became longer in the *scl28* mutant (Fig. 3a, b), which is consistent with the existence of an additional cell size checkpoint at G1/S that compensates for shortened G2 (Jones et al., 2017). This suggests that SCL28 actively inhibits progression through G2 and prevents entry into mitosis. In agreement with the role of SCL28 in G2 duration of the mitotic cell cycle, our ploidy analysis of developing leaves showed that SCL28 also affects endoreplication (Fig. 3c, d, and Supplementary Fig. 4). During leaf development, *SCL28*^OE^ plants initiated earlier endoreplication, indicated by 8C cell emergence as early as eight days after sowing (DAS) and thereafter consistently showing higher cellular ploidy levels compared with WT. Elevated ploidy levels were associated with dramatically increased cell size in *SCL28*^OE^ leaves as shown earlier (Fig. 2b, d). In the *scl28* mutant, ploidy levels were not affected during early stages of leaf development and only showed a modest decrease at 20 DAS. During the earliest stage (8 DAS) before onset of endoreplication, we observed a smaller proportion of 4C cells in *scl28* compared with WT (Fig. 3c), which supports the cell cycle analysis, showing reduced G2 duration (Fig. 3a). In summary, our data suggest that SCL28 has a role to inhibit the G2 to M phase transition.

**Fig. 3.**
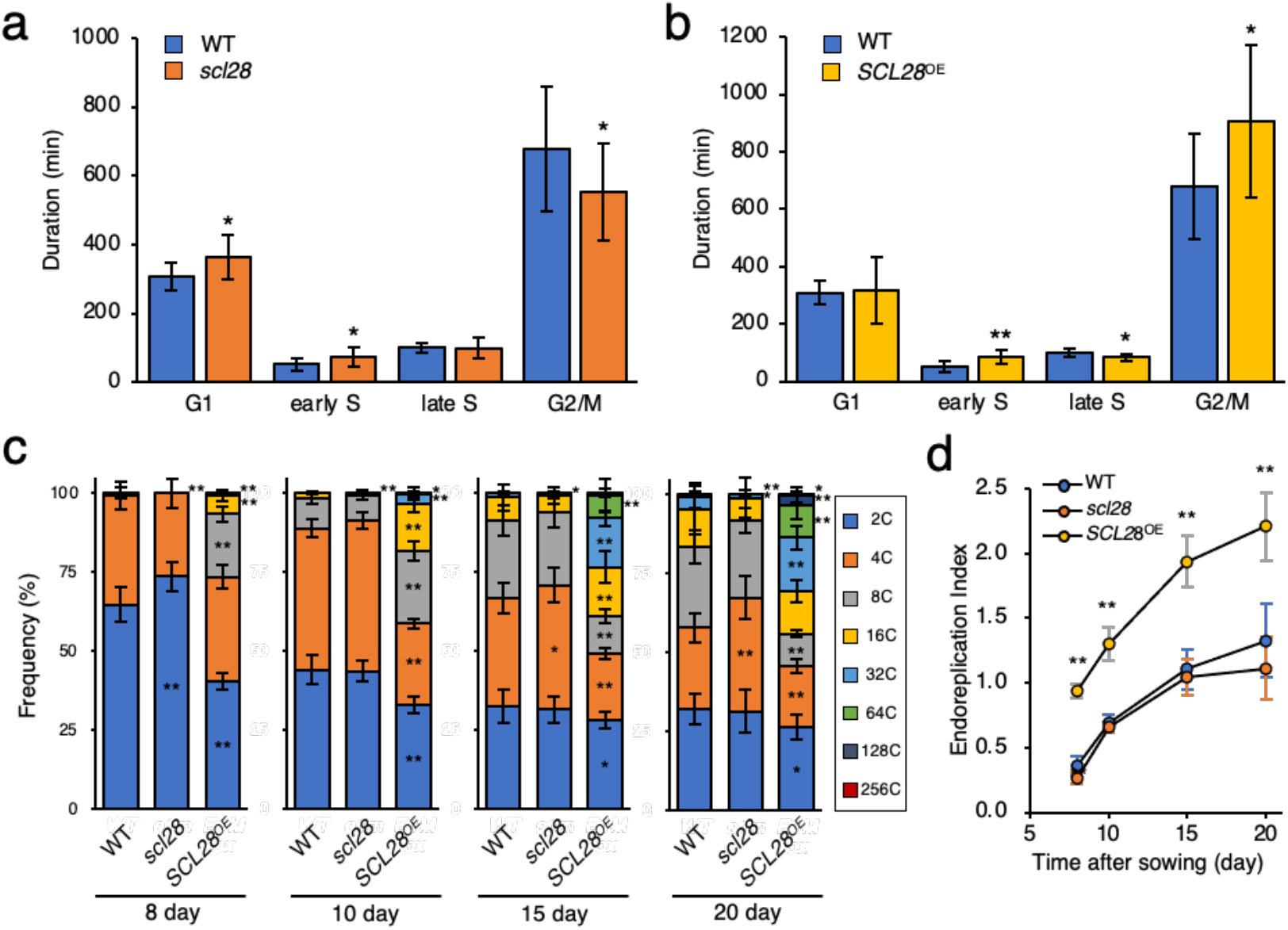
SCL28 inhibits G2 progression and induces endoreplication. **a** G2 duration is shortened in *scl28*. Cortical cells in root meristems of WT, *scl28*, and *SCL28*^OE^ plants carrying the PCNA-GFP marker were analyzed by live-cell imaging for measuring length of G1, early S, late S, and G2/M phases in the cell cycle. Data are shown as mean ± SD (*n* ≥ 10). Statistical significance compared with WT was determined using Student’s t-test. \**P* < 0.05. **b** Prolonged G2 duration in *SCL28*^OE^ plants. Cell cycle analysis in the root meristem was performed as in (**a**). Data are shown as mean ± SD (*n* ≥ 10). Statistical significance compared with WT was determined using Student’s t-test. \**P* < 0.05, \*\**P* < 0.01. **c** Ploidy analysis of *scl28* and *SCL28*^OE^ plants. First leaf pairs of WT, *scl28*, and *SCL28*^OE^ plants were subjected to flow cytometric analysis to determine ploidy distribution during leaf development. Data are shown as mean ± SD (*n* = 10). Statistical significance compared with WT was determined using Student’s t-test. \**P* < 0.05, \*\**P* < 0.01. **d** Time-course change in cellular ploidy levels during leaf development. Data presented in (**c**) were used for calculating endoreplication index, which represents mean number of endoreplication cycles per leaf cells. Data are shown as mean ± SD (*n* = 10). Statistical significance compared with WT was determined using Student’s t-test. \*\**P* < 0.01.

### SCL28 acts together with AtSMOS1 as a heterodimer

Phylogenetic analysis of GRAS family proteins indicated that SCL28 constitutes a unique clade together with SMOS2 in rice (Supplementary Fig. 5a). Rice *smos2* was initially identified as a mutant showing reduced organ size with component cells smaller than those in WT (Hirano et al., 2017). Another rice mutant, *smos1*, with a phenotype similar to that in *smos2*, has loss-of-function mutation in a unique gene encoding an AP2-type transcription factor (Aya et al., 2014). Because a physical interaction between SMOS1 and SOMS2 has been reported (Hirano et al., 2017), we postulated that SCL28 may act through interaction with an Arabidopsis protein orthologous to SMOS1. When AP2-type transcription factors were phylogenetically analyzed, we found At2g41710 to be the likely Arabidopsis ortholog of SMOS1 (Supplementary Fig. 5b). To test the physical interaction between SCL28 and At2g41710, we performed yeast two-hybrid and bimolecular fluorescence complementation (BiFC) assays to obtain clear results showing that they indeed interact in yeast and in plant cells, respectively (Fig. 4a, b). We concluded that the observed protein-protein interaction is evolutionarily related to the SMOS1-SMOS2 interaction in rice and named At2g41710 as AtSMOS1.

**Fig. 4.**
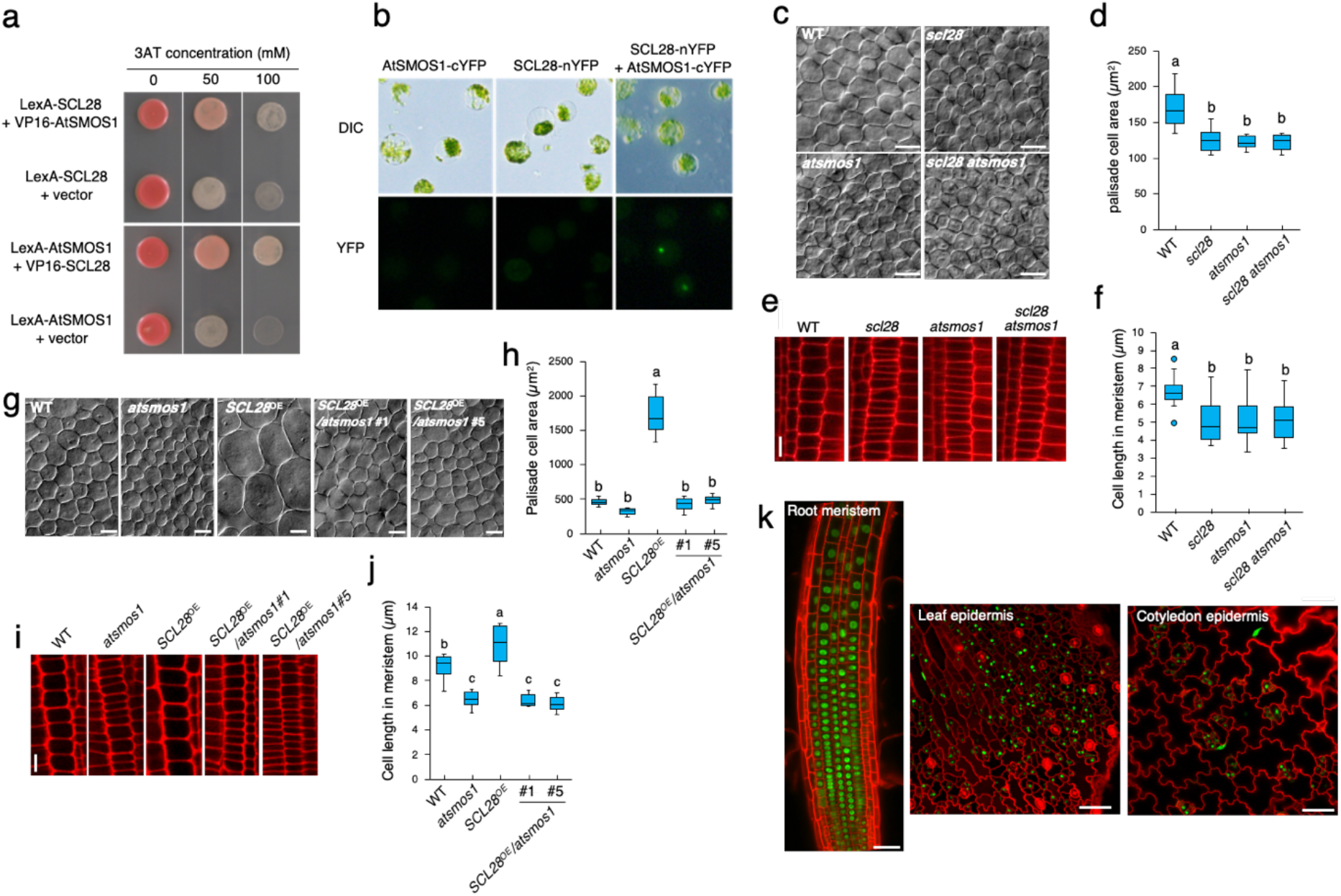
Formation of active heterodimer between SCL28 and AtSMOS1. **a** SCL28-AtSMOS1 interaction in yeast two-hybrid assay. Yeast strains carrying two indicated constructs were grown on synthetic medium lacking His and containing indicated concentration of 3-amino-1,2,4-triazole (3-AT). **b** SCL28-AtSMOS1 interaction in bimolecular fluorescence complementation (BiFC) analysis. Protoplasts prepared from leaf mesophyll cells were transfected with the indicated plasmid constructs. Upper panels, DIC observation; lower panels, epifluorescence microscopic observation. **c** Leaf palisade cells were observed by DIC microscopy using first leaf pairs from WT plants, and *scl28*, *atsmos1*, and double *scl28 atsmos1* mutants at 10 DAS. Scale bar indicates 20 *μ*m. **d** Quantification of palisade cell area in first leaf pairs from plants with indicated genotypes. Data are shown as boxplot (midline = median, box = IQR, whiskers = 1.5 × IQR). Different letters above boxplots indicate significant differences based on one-way ANOVA and Tukey’s test, *P* < 0.05 (*n* = 10). **e** Cortical cell files were observed by CLSM using PI-stained meristems of primary roots from WT plants, and *scl28*, *atsmos1*, and double *scl28 atsmos1* mutants at 5 DAS. Scale bars indicate 10 *μ*m. **f** Quantification of cortical cell length in root meristems from plants with indicated genotypes. Data are shown as boxplots as in (**d**). Different letters above boxplots indicate significant differences as in (**d**) (*n* = 20). **g** DIC observation of palisade cells from first leaf pairs of WT, *atsmos1*, *SCL28*^OE^ plants, and those possessing *atsmos1* and *SCL28*^OE^ in combination. Plants at 15 DAS were used. Scale bar indicates 20 *μ*m. **h** Quantification of palisade cell area in first leaf pairs from the plants with indicated genotypes. Data are shown as boxplots as in (**d**). Different letters above the boxplots indicate significant differences as in (**d**) (*n* = 9). **i** CLSM observation of PI-stained root meristems from WT, *atsmos1*, *SCL28*^OE^ plants, and those possessing *atsmos1* and *SCL28*^OE^ in combination. Plants at 8 DAS were used for observation of cortical cell files. Scale bar indicates 10 *μ*m. **j** Quantification of cortical cell length in root meristems of plants with indicated genotypes. Data are shown as boxplots as in (**d**). Different letters above boxplots indicate significant differences as in (**d**) (*n* ≥ 5). **k** Accumulation patterns of AtSMOS1-GFP protein. Root meristem, and epidermis of leaf and cotyledon from plants at 6 DAS carrying proAtSMOS1::AtSMOS1-GFP were analyzed by CLSM after counterstaining of the cell wall with PI. Scale bars indicate 50 *μ*m.

To examine AtSMOS1 function, we analyzed a T-DNA insertion mutant for this gene (Supplementary Fig. 6) and found a small cell size phenotype that closely resembled that of *scl28* (Fig. 4c–f). However, the overall stature of *atsmos1* was largely indistinguishable from WT plants, as is the case for *scl28* (Supplementary Fig. 7a). To analyze epistasis between *scl28* and *atsmos1*, we performed genetic analysis by quantitatively comparing phenotypes of *scl28*, *atsmos1* and *scl28 atsmos1* double mutants. These single and double mutants showed essentially equivalent phenotypes in terms of palisade cell size (Fig. 4c, d) and cell size in the root meristem (Fig. 4e, f). This indicates that, as with SCL28, AtSMOS1 also impacts cell size. The lack of additive effect between *atsmos1* and *scl28* suggests that these proteins may act in the same pathway, which is consistent with the interaction between these proteins. We then tested whether the enlarged cells in *SCL28*^OE^ relies on the presence of AtSMOS1, and indeed in the *atsmos1* mutant background, the strong effects of *SCL28*^OE^ on cell size of leaf palisades (Fig. 4g, h) and root meristem cells (Fig. 4i, j), as well as whole plant growth, completely diminished (Supplementary Fig. 7b). Collectively, these observations strongly suggest that SCL28 and AtSMOS1 function cooperatively to regulate cell size in different plant organs.

To analyze AtSMOS1 expression, we generated plants expressing AtSMOS1-GFP driven by its own promoter. These transgenic plants showed nuclear localization of AtSMOS1-GFP in various tissues and organs such as root meristem, developing leaves, and cotyledons (Fig. 4k). However, unlike SCL28, AtSMOS1-GFP was uniformly expressed in meristematic cells, suggesting a cell cycle-independent AtSMOS1 expression. Nonetheless, cells expressing AtSMOS1 and SCL28 were overlapping, supporting the idea that they interact to form a heterodimer *in vivo*. We also noted an E2F binding element in the *AtSMOS1* promoter, which binds RBR in published ChIP-Seq data (Bouyer et al., 2018). This highlights the possibility that, while *SCL28* shows mitotic control through the MYB3Rs, *AtSMOS1* may be regulated by the RBR-E2F pathway.

### Downstream targets of SCL28

Studies of *smos1* and *smos2* rice mutants suggest a role in post-mitotic cell expansion through the regulation of the *Oryza sativa PHOSPAHATE INDUCED1* (*OsPHI-1*) gene (Hirano et al., 2017). Because we observed SCL28 expression specific to meristematic cells, we postulated that SCL28 should largely influence actively-proliferating cells before the onset of cell expansion. To identify the downstream targets of SCL28, we analyzed genome-wide gene expression changes in *scl28*, *SCL28*^OE^, and *atsmos1* by conducting RNA-sequencing (RNA-Seq) and microarray experiments. Considering the epistatic phenotypes among *scl28*, *SCL28*^OE^, and *atsmos1* mutants, the critical downstream genes of SCL28 should be affected in *scl28* and *atsmos1* in a similar manner, as well as in *SCL28*^OE^ in an opposing manner. To reveal the genes satisfying these expression criteria, we conducted an overlapping analysis of differentially expressed genes in *scl28*, *atsmos1*, and *SCL28*^OE^, and identified 21 genes that are significantly (adj *P*-value less than 0.05) downregulated in both *scl28* and *atsoms1*, and upregulated in *SCL28*^OE^ (Fig. 5a and Supplementary Fig. 8a), as well as 26 genes affected in an opposing way within each line (Fig. 5b and Supplementary Fig. 8b). Among these genes, we focused on *SIAMESE-RELATED2* (*SMR2*), encoding a member of SMR family proteins, as proteins of this family represent plant-specific CDK inhibitors and some members, such as SIM and SMR1, positively affect cell size and negatively influence cell division, reflecting the observed effect of SCL28 (Roeder et al., 2010; Kumar et al., 2015). According to RNA-Seq and microarray data, in addition to *SMR2*, we found other *SMR* family members as candidates of downstream genes, with decreased expression levels in both *scl28* and *atsmos1* and increased levels in *SCL28*^OE^. To further analyze *SMR* family genes, we conducted quantitative reverse-transcription PCR (qRT-PCR) analysis to examine expression of all members of this family in *scl28, atsmos1*, and *SCL28*^OE^, and identified at least seven members (*SMR1*, *SMR2*, *SMR6*, *SMR8*, *SMR9*, *SMR13*, and *SMR14*) out of all 17 as potential target genes of SCL28 and AtSMOS1 (Fig. 5c-e). Consistent with this conclusion, we also observed reduced expression of proSMR2::SMR-GFP and proSMR13::SMR13-GFP in root meristems in *scl28* or *atsoms1* mutant backgrounds compared with WT (Supplementary Fig. 9).

**Fig. 5.**
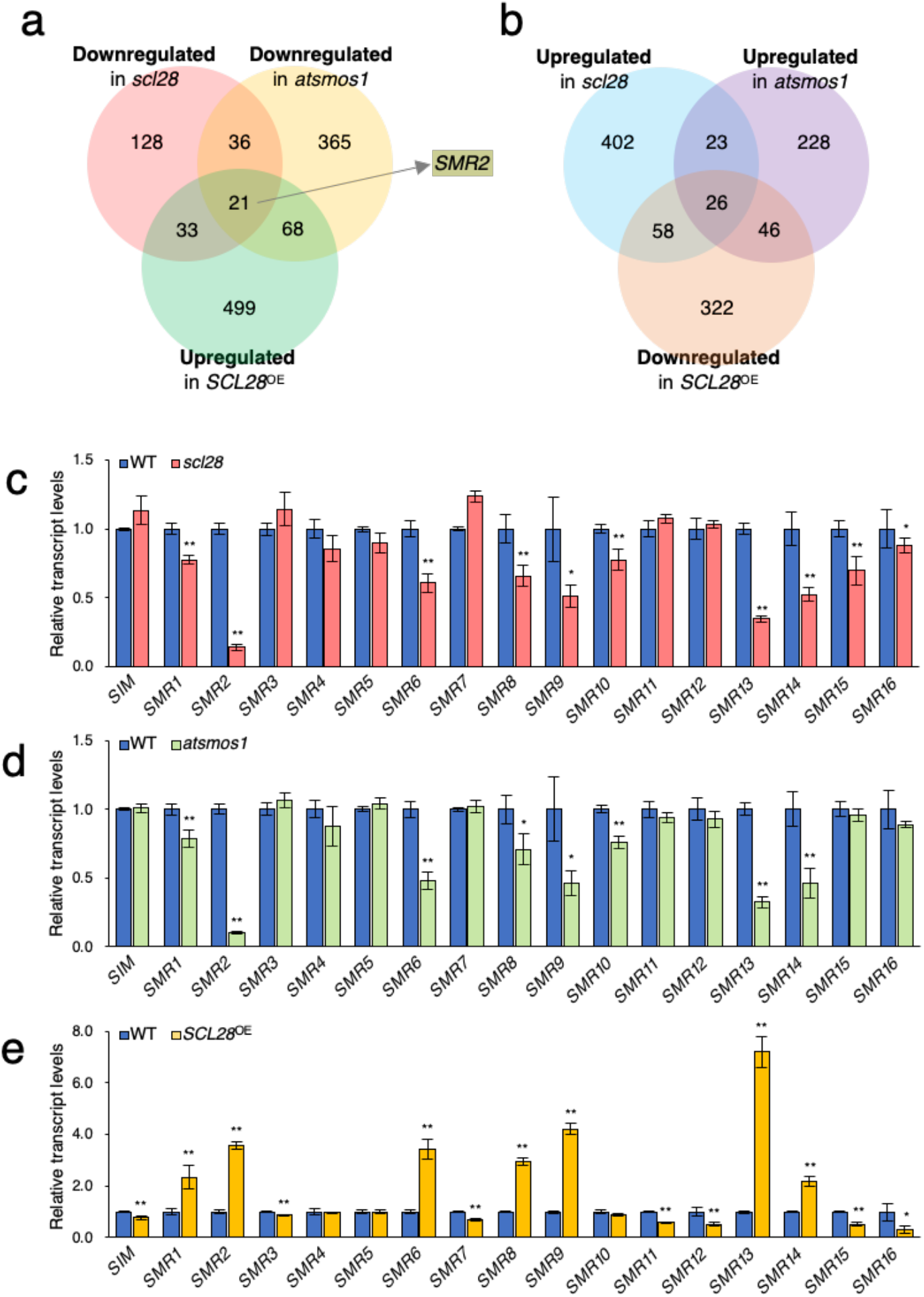
SCL28 and AtSMOS1 activate transcription of *SMR* genes. **a** Venn diagram representing overlap between genes downregulated in *scl28* and *atsmos1*, and upregulated in *SCL28*^OE^ plants. The *SMR2* gene was found in the overlap of these three categories. **b** Venn diagram representing overlap between genes upregulated in *scl28* and *atsmos1*, and downregulated in *SCL28*^OE^ plants. **c–d** Transcript levels of *SMR* genes in *scl28* (**c**), *atsmos1* (**d**), and *SCL28*^OE^ (**e**) plants at 8 DAS. qRT-PCR analysis of all 17 genes in the SMR family was performed to compare expression levels with WT plants. Data are shown as mean ± SD (*n* = 3). Statistical significance compared with WT was determined using Student’s t-test. \**P* < 0.05, \*\**P* < 0.01.

KIP-RELATED PROTEINs (KRPs) constitute another family of CDK inhibitors in addition to SMRs in plants. The general view is that KRP proteins primarily connect with CDKA to negatively regulate G1/S (Verkest et al., 2005), while SMR proteins inhibit G2/M by associating with both CDKA and CDKB1 (Kumar et al., 2015; Wang et al., 2020). There are seven KRP members encoded in the Arabidopsis genome, none of which showed significant change in expression for all *scl28*, *atsmos1*, and *SCL28*^OE^ plants in our transcriptome analysis (Supplementary Fig. 10). Therefore, SCL28 and AtSMOS1 affect expression of SMR members, but not those of the KRP family, supporting our results that SCL28 preferentially affects G2 duration in meristematic cells.

### Genome-wide mapping of SCL28 and AtSMOS1 binding sites

In order to define direct targets for SCL28 and AtSMOS1, combined with transcriptome analysis, we performed ChIP-Seq assays using proRPS5A::SCL28-GFP and proAtSMOS1::AtSMOS1-GFP lines to identify their binding genomic loci. Consistent with our emphasis on downregulation of *SMR* genes, which suggests that SCL28 and AtSMOS1 act as transcriptional activators, the majority of binding sites are located in promoter regions proximal to transcription start sites (TSS) (Supplementary Fig. 11) and located within nucleosome-free and highly accessible chromatin regions (Supplementary Fig. 12a-d). Their enrichment levels at TSS correlated well with mRNA levels of corresponding genes, suggesting a positive correlation between their binding and transcriptional activity (Supplementary Fig. 12e, f).

The ChIP-Seq data analysis revealed 463 and 4,287 genes as targets of SCL28 and AtSMOS1, respectively. Comparing these target genes revealed the presence of a significant overlap, a set of 308 common targets that accounts for 66% of SCL28 targets (Fig. 6a). On one hand, this result confirms our genetic analysis, suggesting that SCL28 function largely relies on the interaction with AtSMOS1. On the other hand, AtSMOS1 may show a wider range of functions both dependent on and independent of SCL28. We found only limited overlap between common targets identified by ChIP-Seq and those that show regulated expression in transcriptome analysis of *scl28*, *atsmos1* and *SCL28*^OE^ lines (Fig. 6b). Gene ontology (GO) enrichment analysis of the 308 common targets revealed overrepresented GO terms related to cell cycle, such as “regulation of mitotic nuclear division” and “regulation of DNA endoreduplication” (Supplementary Fig. 13). This overrepresentation of cell cycle-related GO terms largely relies on the presence of five common genes that belong to such GO categories. All these genes were found to be members of *SMR* family genes, *SMR2*, *SMR4*, *SMR6*, *SMR8*, and *SMR9*, all of which, except *SMR4*, showed significant downregulation in both *scl28* and *atsmos1* and upregulation in *SCL28*^OE^ in our qRT-PCR experiments described above (see Fig. 5). Though three *SMR* genes—*SMR1*, *SMR13*, and *SMR14*—with significant expression changes in all lines failed to fulfill the criteria of ChIP-Seq data analysis, visual inspection revealed recognizable peaks for both SCL28 and AtSMOS1 in ChIP-Seq profiles, suggesting these *SMRs* are also direct common targets (Fig. 6c and Supplementary Fig. 14). The ChIP-Seq peaks were observed at either 5’ (*SMR1, SMR2*, and *SMR4*), 3’ (*SMR6*, *SMR9*, and *SMR14*) or both 5’ and 3’ (*SMR8* and *SMR13*) regions of the target *SMR* loci. In most cases, we found that the positions of ChIP-Seq peaks adjacent to *SMR* loci coincided for SCL28 and AtSMOS1, suggesting that they associate with the same sites.

**Fig. 6.**
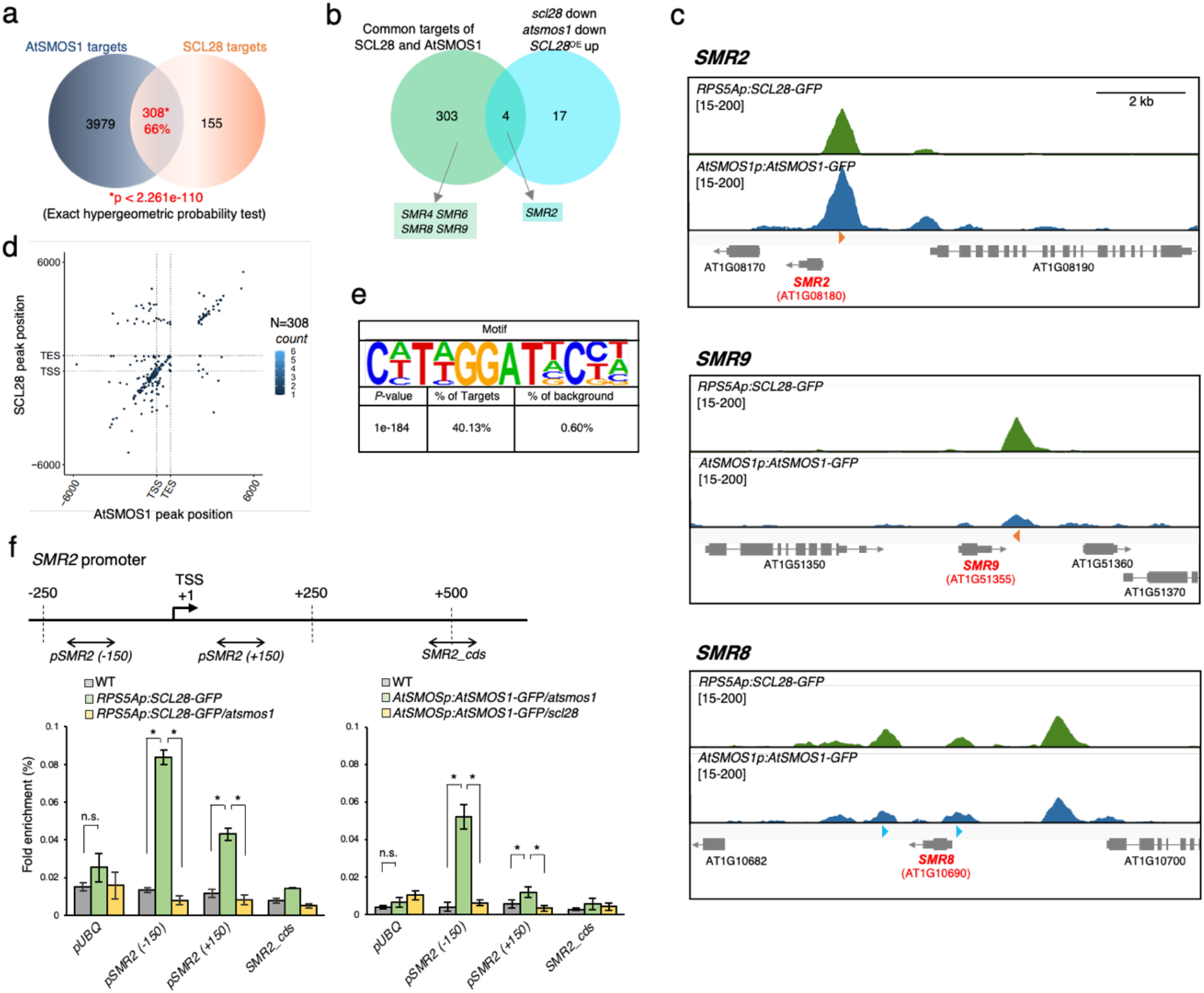
Binding of active SCL28-AtSMOS1 heterodimer to target *SMR* genes. **a** Venn diagram showing significant overlap between genes bound by AtSMOS1 and SCL28. **b** Venn diagram showing limited overlap between common target genes bound by SCL28 and AtSMOS1 and those downregulated in both *scl28* and *atsmos1* and upregulated in *SCL28*^OE^. SMR genes found in overlapping and non-overlapping regions are also shown. **c** ChIP-Seq profiles for SCL28 and AtSMOS1 around *SMR2*, *SMR8*, and *SMR9* loci. Orange arrowheads indicate DNA motifs perfectly matching C(a/t)T(a/t)GGATNC(c/t)(a/t), identified as an enriched motif in the SCL28/AtSMOS1 common targets, whereas blue arrowheads indicate motifs matching the enriched motif with one base mismatch (see Fig. 6e). **d** Density plot showing overlap of SCL28 and AtSMOS1 using hexagonal binning routine. Each dot represents the distance from the peak midpoint to the nearest gene. The *y*-axis shows location of the AtSMOS1 peak midpoint compared with gene position, while the *x*-axis indicates location of SCL28 peak midpoint compared with nearest gene. The large number of dots occurs along the positive correlation line, showing the co-occurrence pattern of SCL28 and AtSMOS1. **e** DNA motif overrepresented in common targets for SCL28 and AtSMOS1. HOMER motif search identified a specific common SCL28/AtSMOS1 binding motif with a *P*-value of 1e-184, which was found in 40.13% of common targets. **f** ChIP-qPCR analysis was performed on *SMR2* loci using plants carrying proRPS5A::SCL28-GFP under WT and *atsmos1* backgrounds, and those carrying proAtSMOS1::AtSMOS1-GFP under *scl28* and *atsmos1* backgrounds. Amplified regions (−150, +150, and cds) in ChIP-qPCR are shown by doubleheaded arrows. *UBQ* locus was analyzed in the same way as a negative control without binding. Data are shown as mean ± SD (*n* = 3). Statistical significance was determined using Student’s t-test. \**P* < 0.05. n.s., not significant.

To find common binding sites and sequence motifs for SCL28 and AtSMOS1, we compared the precise positions of ChIP-Seq peaks along the gene structure. Dot-plot analysis of the common targets showed frequent clustering of SCL28 and AtSMOS1 peaks at the same positions relative to TSS and transcription end site (TES; Fig. 6d). Similar analysis at a genome-wide scale confirmed that SCL28 frequently targets the same sites as AtSMOS1 around the TSSs (Supplementary Fig. 15a). Within the common peaks for SCL28 and AtSMOS1, a DNA motif of C(a/t)T(a/t)GGATNC(c/t)(a/t) could be identified as an overrepresented *cis* element. This motif is indeed present among more than 40% of the common targets, but is much less frequent among targets of either SCL28 or AtSMOS1, which contain quite different *cis* elements based on sequence overrepresentation analysis (Supplementary Fig. 15b, c). This suggests that the formation of SCL28-AtSMOS1 heterodimer generates a specific sequence preference for DNA binding to regulate transcription of a specific set of targets. This or closely related motifs are present in all seven *SMR* target loci at positions frequently coinciding with the peak summits of SCL28 and AtSMOS1 (Fig. 6c and Supplementary Fig. 14), suggesting that SCL28 and AtSMOS1 recognize this motif and bind to their targets as a heterodimer.

We then tested whether chromatin association of SCL28 depended on AtSMOS1 and vice-versa by conducting ChIP-qPCR experiments in *scl28* and *atsmos1* backgrounds (Fig. 6f). As expected, binding of both SCL28 on *SMR2* promoter in the *atsmos1* and AtSMOS1 binding in the *scl28* mutant backgrounds were abolished. Collectively, our data supports SCL28 and AtSMOS1 functional heterodimer binding to specific sets of *SMR* target genes to activate transcription and negatively regulate cell cycle progression at the G2 to M cell cycle transition.

### *SMRs* are critical downstream effectors for SCL28 and AtSMOS1

If *SMR* genes directly bound by SCL28 and AtSMOS1 are crucial for mediating their effects on cellular phenotype, transcript levels of those *SMR* genes should be altered accordingly with cell size phenotype observed in our epistasis analysis (see Fig. 4). Confirming this idea, downregulation of target *SMR* genes—*SMR2*, *SMR9* and *SMR13*—were quantitatively equivalent in *scl28*, *atsmos1*, and double *scl28 atsmos1* mutants (Fig. 7a). In addition, strong activating effects of *SCL28*^OE^ on target transcription was totally abolished under *atsmos1* mutant background, providing the molecular basis for AtSMOS1-dependent action of SCL28 on cell size (Fig. 7b). To further elucidate that AtSMOS1 acts together with SCL28 on *SMR* genes, we also analyzed the proSMR2::LUC reporter co-transfected in protoplasts with pro35S::SCL28 and/or pro35S::AtSMOS1 constructs. The most prominent and significant activation of luciferase was only observed when both plasmids expressing SCL28 and AtSMOS1 were simultaneously transfected, supporting the conclusion that SCL28 and AtSMOS1 fulfill their function as a transcriptional activator by forming a heterodimer (Fig. 7c).

**Fig. 7.**
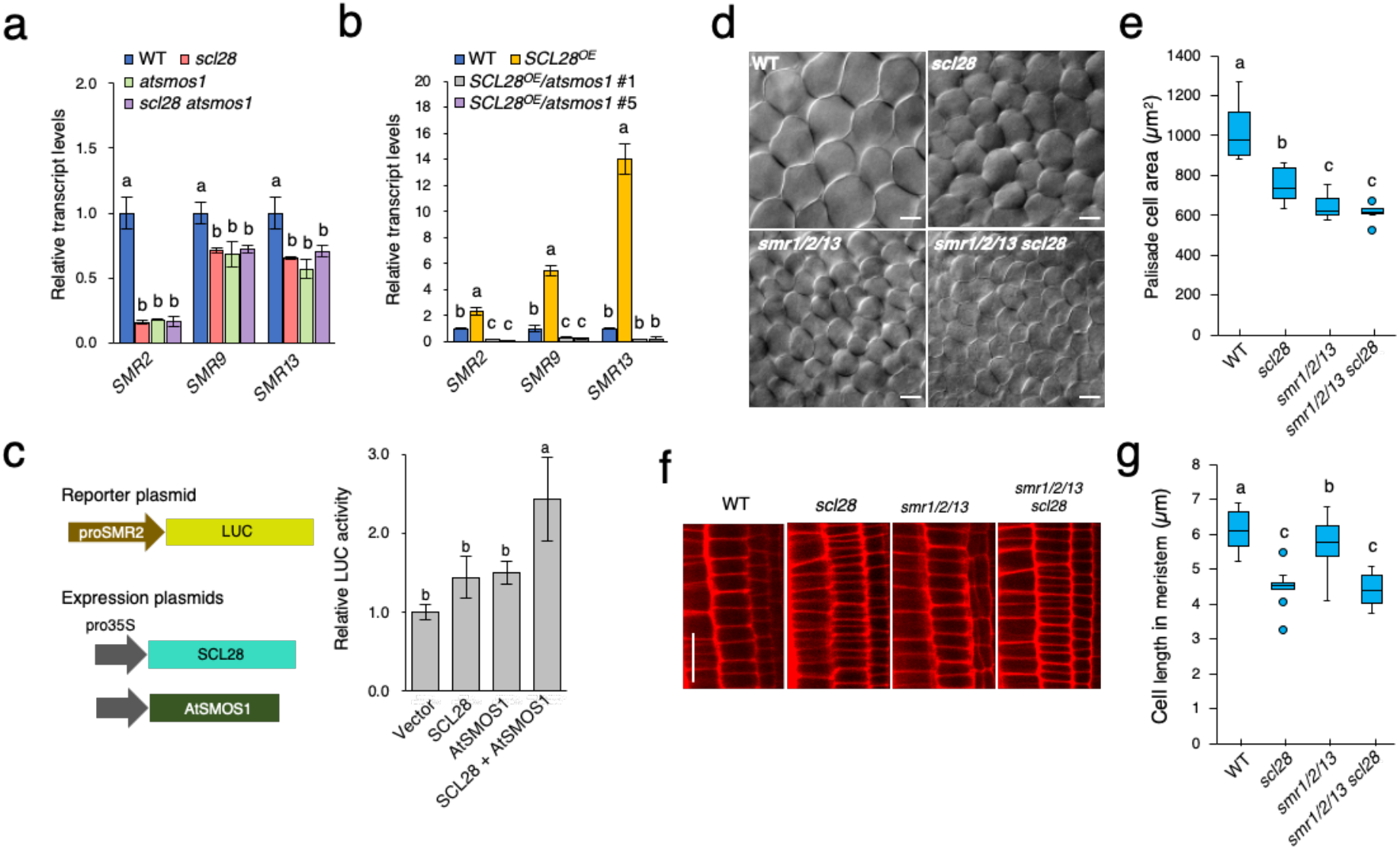
Cell size control by SCL28-AtSMOS1 is mediated by *SMRs*. **a** Double *scl28 atsmos1* mutation downregulates *SMR* genes, equivalently to each single mutation. qRT-PCR analysis of *SMR2*, *SMR9* and *SMR13* transcripts was performed for comparing expression levels in WT, *scl28*, *atsmos1*, and *scl28 atsmos1* plants. Data are shown as mean ± SD (*n* = 3). Statistical analysis was performed for each *SMR* gene based on one-way ANOVA and Tukey’s test, and significant differences (*P* < 0.05) were shown in different letters above the bars. **b** Upregulation of *SMR* genes in *SCL28*^OE^ requires presence of AtSMOS1. qRT-PCR analysis was performed for comparing expression levels between *atsmos1*, *SCL28*^OE^ plants, and those carrying *atsmos1* and *SCL28*^OE^ in combination. Two independent lines were used for analyzing *SCL28*^OE^/ *atsmos1*. Data were statistically analyzed as in (**a**). **c** Co-expression of SCL28 and AtSMOS1 activates the *SMR2* promoter in the protoplast transient assay. proSMR2::LUC reporter plasmid was transfected into protoplasts prepared from T87 cells together with expression plasmids of SCL28 and AtSMOS1 as indicated. Schematic presentation of reporter and expression plasmids as shown on the left. Data were statistically analyzed as in (**a**) **d** DIC observations of leaf palisade cells in WT, *scl28*, *smr1*/*2*/*13*, and *smr1*/*2*/*13 scl28* plants at 22 DAS. **e** Quantification of palisade cell area in the first leaf pairs from plants with indicated genotypes. Quantified data are shown as boxplot (midline = median, box = IQR, whiskers =1.5 × IQR). Different letters above the boxplots indicate significant differences based on one-way ANOVA and Tukey’s test. *P* < 0.05 (*n* = 10). **f** CLSM observation of cortical cell files in root meristems from WT, *scl28*, *smr1*/*2*,/*3*, and *smr1*/*2*/*13 scl28* plants at 7 DAS. Scale bar indicates 20 *μ*m. **g** Quantification of cortical cell length in root meristems from plants with indicated genotypes. Quantified data are shown as boxplots as in (e). Different letters above boxplots indicate significant differences as in (e) (*n* = 10).

To directly address whether these *SMR* genes are critical for SCL28 function, we performed genetic analysis focusing on *SMR1*, *SMR2* and *SMR13*, which are significantly downregulated in *scl28* and upregulated in *SCL28*^OE^. For each *smr* mutant, we could not detect any abnormalities in cell size (Supplementary Fig. 16). However, when these mutations were combined in a *smr1*/*2*/*13* triple mutant, there was a significant reduction in cell size in multiple tissues, including leaf palisades (Fig. 7d, e) and root meristem (Fig. 7f, g), suggesting functional redundancy among *SMR* genes. To address the link between SCL28 and SMRs, we studied the cell size phenotype when these mutations were combined. In leaf palisade cells, the triple *smr1*/*2*/*13* mutant showed stronger reduction in cell size than the *scl28* mutant, which did not further reduce when all mutations combined in the *smr1*/*2*/*13 scl28* line, supporting the idea that cell size regulation by SCL28 is mediated by SMR1/2/13 (Fig. 7d, e). Conversely, *scl28* mutation still significantly decreased cell size in the root meristem when combined with *smr1*/*2*/*13* (Fig. 7f, g). Therefore, *SMR1*/*2*/*13* contribution downstream of SCL28 appears to be larger in palisade tissue than root meristem, where additional SCL28-regulated *SMRs* may play a role in cell size control. These tissue-specific differences indicate the contribution of developmental regulation that positions different sets of *SMRs* downstream of SCL28.

### Dose-dependent control of cell size and cell number by SCL28 to maintain organ size homeostasis

In the *scl28* mutant, leaf growth is essentially normal, but the constituent cells became small and more numerous (Fig. 2d, e, g), suggesting that SCL28 is pivotal to maintain organ size homeostasis by regulating the balance between cell size and cell number. For a regulator designed to tune the balance between cellular parameters, one assumes that it acts dose dependently. To explore whether or not SCL28 levels quantitatively affect cell size, we analyzed plants heterozygous for *scl28* (*scl28*/+ plants). Heterozygous plants showed reduced *SCL28* transcript levels that is approximately half of that in WT (Fig. 8a). This *SCL28* downregulation resulted in a clear reduction of cell size in root meristem and leaf palisade tissue, with an intermediate cell size in heterozygous plants between those in WT and *scl28* homozygous plants (Fig. 8b, c and Supplementary Fig. 17a, b).

**Fig. 8.**
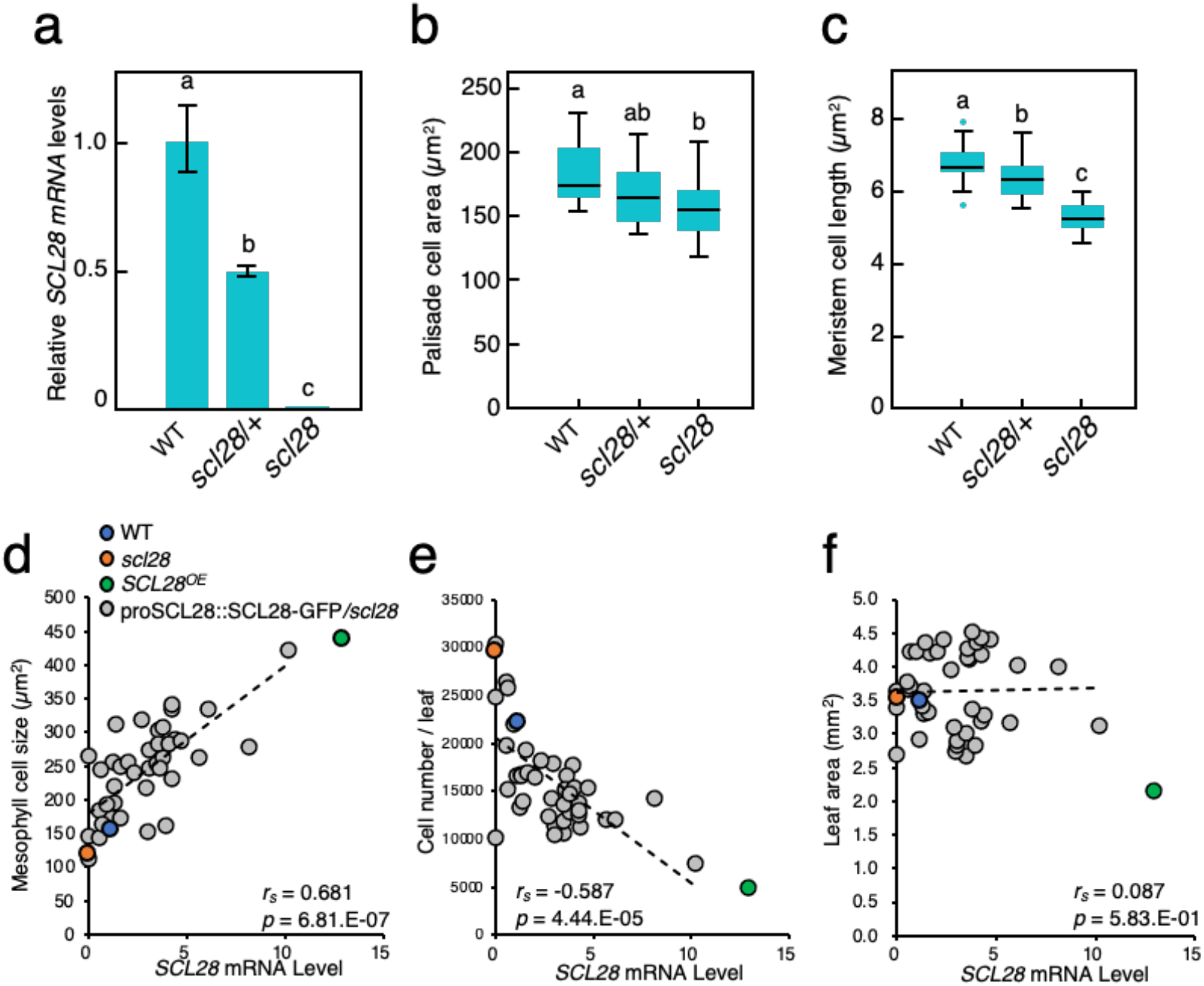
*SCL28* regulates cell size in a dose-dependent manner. **a** Moderate downregulation of *SCL28* transcripts in plants heterozygous for *scl28*. qRT-PCR analysis was performed using WT plants, and those heterozygous (*scl28/+*) and homozygous (*scl28*) for *scl28* at 10 DAS. Data are shown as mean ± SD (*n* = 3). Statistical analysis was performed based on one-way ANOVA and Tukey’s test and significant differences (*P* < 0.05) are shown in different letters above bars. **b** Quantification of palisade cell area in *scl28*, *scl28*/+ and WT plants at 10 DAS. Data are shown as boxplots (midline = median, box =IQR, whiskers = 1.5 × IQR), where different letters above boxplots indicate significant differences based on one-way ANOVA and Tukey’s test, *P* < 0.05 (*n* ≥ 15). **c** Quantification of cortical cell length in root meristems of *scl28*, *scl28*/+ and WT plants at 7 DAS. Data are shown as boxplots as in (**b**). Different letters above the boxplots indicate significant differences as in (**b**) (*n* ≥ 12). **d** Correlation between *SCL28* mRNA levels and palisade cell size analyzed by scatterplots, where each dot corresponds to each individual plant from different transgenic lines carrying proSCL28::SCL28-GFP under *scl28* background. In addition to proSCL28::SCL28-GFP plants (shown in gray dots), WT (blue dot), *scl28* (red dot), and *SCL28*^OE^ plants (green dot) were also analyzed. **e** Correlation between *SCL28* mRNA levels and number of palisade cells per leaf was analyzed by scatterplot as in (**d**). Symbols are same as in (**d**). **f** Correlation between *SCL28* mRNA levels and leaf area was analyzed by scatterplot as in **(d)**. Symbols are same as in **(d)**.

Overexpression of *SCL28* causes a phenotype opposite to *scl28*, dramatically reducing the number of cells that became enlarged. To examine the outcome when SCL28 expression is only moderately increased, we utilized plants from different T2 lines carrying proSCL28::SCL28-GFP in *scl28* mutant background and showing varying levels of SCL28-GFP expression. We quantitatively compared levels of *SCL28* transcript and cellular parameters of palisade tissues in each individual plant (Fig. 8d, e and Supplementary Fig. 17c). This showed that SCL28 expression positively correlates with cell size and negatively with leaf cell number. Both positive and negative correlations were statistically significant. Due to compensatory changes in cell number and size, overall leaf size was not dramatically affected by moderate alteration of SCL28 expression (Fig. 8f). Therefore, we concluded that a finely-graded expression of SCL28 sets the balance between cell size and number without altering organ growth as a whole. This mechanism may enable plants to achieve an optimal balance between cell size and number, possibly depending on developmental status and environmental conditions.

## Discussion

Cell size at division depends on coordination between cell growth and cell cycle progression, which is actively maintained in a cell autonomous manner in both uni- and multicellular organisms (Zatulovskiy and Skotheim, 2020; Jones et al., 2019; D’Ario and Sablowski, 2019). Here, we identified a molecular mechanism for cell size regulation in Arabidopsis that relies on a hierarchical transcriptional activation of CDK inhibitors. In this pathway, SCL28 expression is specifically confined to mitosis through the MSA element in its promoter controlled by MYB3Rs. In turn, SCL28 associates with AtSMOS1, a transcription factor with cell cycle-independent expression, and this dimer defines the binding site present in a set of *SMR* genes encoding CDK inhibitors to activate their expression and thereby act as a brake against the cell cycle engine predominantly driven by CDK activity (Fig. 9). The SCL28-AtSMOS1-SMR axis uniquely affects only cell cycle progression and cell doubling time, but not the exit from cell proliferation. Therefore, it acts to set the balance between cell number and size during organ development without significantly impacting final organ size. Whether SCL28 is part of a cell size sensor and how it is coupled to the measurement of cell growth remain to be determined.

**Fig. 9.**
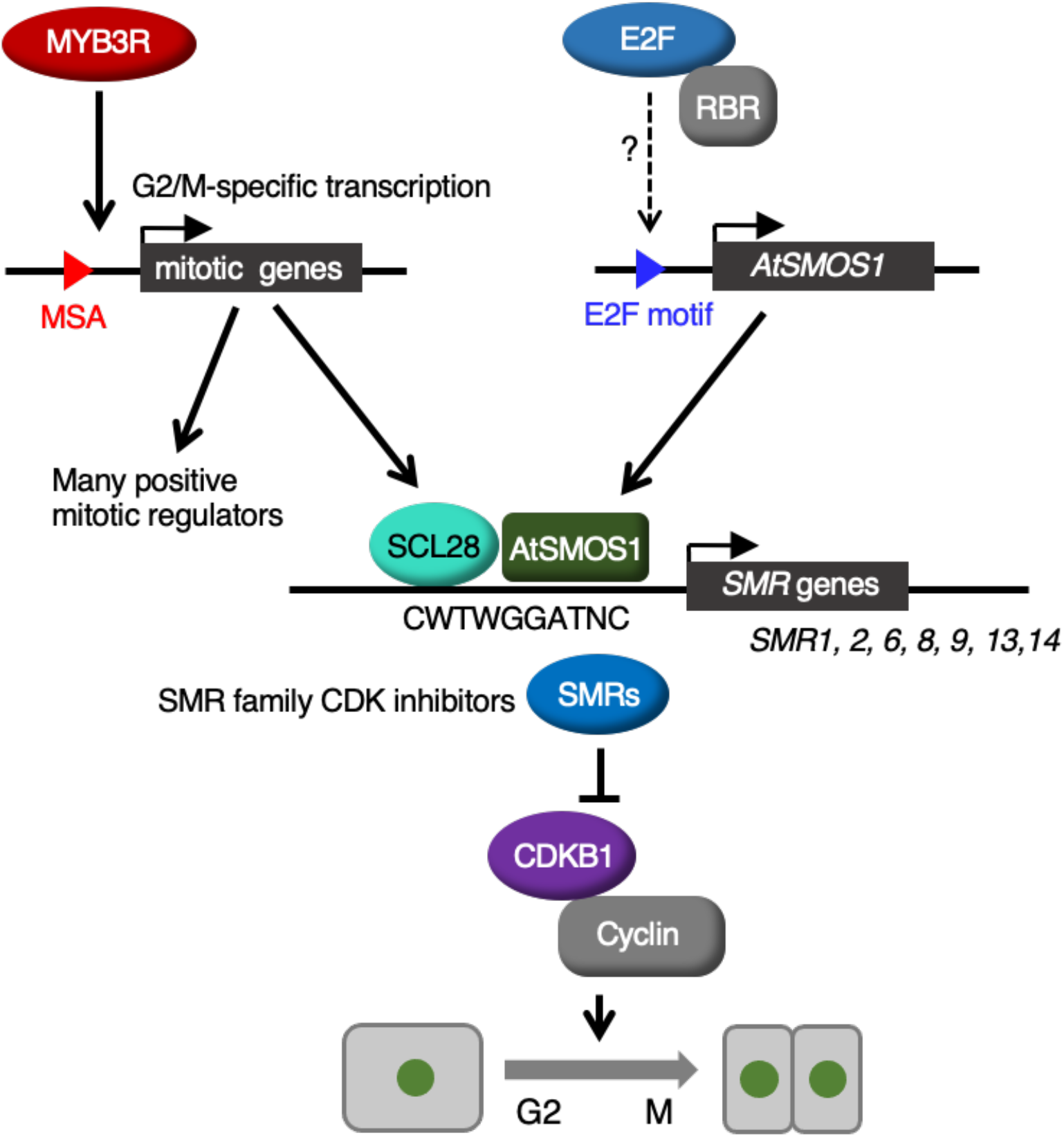
Schematic model of SCL28-dependent cell size control. MYB3R transcription factors regulate a suite of G2/M-specific genes, including many positive mitotic regulators. However, *SCL28*, similarly regulated by MYB3Rs acts as a negative regulator of the cell cycle together with *AtSMOS1*, which may be under the control of the E2F-RBR pathway. Formation of the SCL28-AtSMOS1 heterodimer generates specific binding sequences, enabling direct transcriptional activation of SMR family genes. An increasing amount of SMR proteins, in turn, inhibits CDK activity, thereby negatively influencing cell cycle progression during G2. By controlling the activity of SCL28-AtSMOS1 heterodimer, G2 duration in the cell cycle can be finely tuned to maintain a proper balance between cell size and cell number in developing plant organs.

We provide multiple lines of evidence supporting the conclusion that SCL28 acts as a brake on the G2/M transition to set cell size: 1) measurements of cell cycle phases by following PCNA-GFP using live-cell imaging showed that *scl28* mutant and *SCL28*^OE^ shortened or lengthened the G2 phase, respectively; 2) kinematic analysis of cellular parameters during leaf development of *scl28* and *SCL28*^OE^ showed a clear alteration in cell doubling time without any effect on cell cycle exit; 3) the molecular mechanisms identified here are fully consistent with the cell cycle inhibitory function, SCL28, forming a dimer with AtSMOS1 and directly activating the transcription of *SMR* genes whose products inhibit CDK activity; 4) in support of this, the *scl28* mutant phenocopied and interacted in an epistatic fashion with the triple *smr1*/*2*/*13* mutant; and 5) SCL28 regulates the number of endoreplication cycles, which is typically enhanced by G2 inhibition. Previous reports suggest that SCL28 plays a positive role in root growth and promotes progression through the G2/M cell cycle progression (Choe et al., 2017; Goldy et al., 2021). These conclusions are inconsistent with our data and the small cell size phenotype observed in both studies, which may originate from using different methods relying on fixed samples.

Another example of a transcription factor inhibiting cell cycle at G2/M is a NAC-type transcription factor, SUPPRESSOR OF GAMMA RESPONSE 1 (SOG1), acting as a central regulator of DNA damage response in Arabidopsis. Similar to SCL28, SOG1 transcriptionally activates *SMR* family genes to inhibit cell cycle progression upon DNA damage (Yi et al., 2014), but its targets, *SMR5* and *SMR7*, are not regulated by SCL28, which instead directly regulates seven other *SMR* genes. Notably, in contrast with SOG1, which is activated specifically upon DNA damage, SCL28 negatively regulates cell cycle, thereby affecting cell size during normal plant development. The target of rapamycin (TOR) signaling pathway is a primary regulator of cell growth and proliferation and is involved in cell size regulation in yeast (Davie and Petersen, 2012). In Arabidopsis, TOR, acting through YAK1 kinase, regulates the transcription of *SMR* genes and thereby G2/M transition (Barrada et al., 2019). However, the set of YAK1-targeted *SMRs* overlap with those regulated by SOG1 rather than those regulated by SCL28. Therefore, our study places SCL28 as part of a novel pathway for regulating G2/M progression that determines cell size and cell number during plant organ development.

Variability in cell size at birth or stochastic changes in key regulators necessitate mechanisms that achieve cell size homeostasis by linking cell cycle progression to cell growth (D’Ario and Sablowski, 2019). The way cell size variations are corrected has long been debated. The principal possibilities are using (i) a cell cycle timer, (ii) cell size measurements or (iii) correcting cell size by adding a constant increase to daughters regardless of initial sizes (Zatulovskiy and Skotheim, 2020). Comparing predicted outcomes of these theoretical models with time-lapse imaging data on cell size in plant meristematic cells identifies possible co-existence of a mix of these mechanisms that may act at multiple control points of the cell cycle (Willis et al., 2016; Jones et al., 2017). The molecular device for cell size control may rely on cell size-dependent accumulation of an activator or dilution of a cell cycle inhibitor molecule. The ‘inhibitor dilution model’ postulates that cell growth dilutes such inhibitor protein until its concentration meets a threshold, thereby permitting cell division when cells reach an appropriate size (Fantes et al., 1975). This hypothesis has been verified in yeast and mammalian systems (Schmoller et al., 2015; Zatulovskiy et al, 2020). In Arabidopsis, recent studies show segregation of set amounts of chromatin-bound KRP4 CDK inhibitors act as a cell size sensor that is diluted by cell growth and is responsible for cell size-dependent cell cycle regulation at G1/S (D’Ario et al., 2021).

Live-cell imaging of cell cycle markers in Arabidopsis shoot apical meristem demonstrated that both G1/S and G2/M transitions are regulated in a cell size-dependent manner. Accumulation of two waves of CDK activities underpinned by D-type cyclin and mitotic CDKB1, their interactors or downstream effectors were proposed as molecular components for cell size-dependent G1/S and G2/M regulation, respectively (Jones et al., 2017). SCL28 may participate in regulating G2/M transition in this model, in which dilution of an inhibitor rather than accumulation of an activator is the principle behind. How SCL28 is connected to monitoring of cell size remains to be determined. Stochastic variability in the accumulation of key regulators can also set cell size and cell fate differences in a developing organ. An example is ATML1, a transcription factor that is necessary for the patterning of giant cells within the sepal epidermis by overcoming a threshold in G2 cells and activating the downstream *SMR1*, also called LOSS OF GIANT CELLS FROM ORGANS (LGO) (Meyer et al., 2017). Similar to ATML1, SCL28 shows a finely-tuned dose-dependent effect on cell size by accurately distinguishing gene copy numbers of *SCL28* and reflecting its graded overexpression. It is possible that the molecular ruler in an inhibitor dilution model potentially involving SCL28 is set by the synthesis rate limited by gene copy number, as shown for Whi5 in budding yeast (Schmoller et al., 2015).

The human functional orthologue of Whi5 is pRB, which also controls cell size via its synthesis mostly after G1 and subsequent dilution during growth (Zatulovskiy et al, 2020). In Arabidopsis, E2FB can strongly affect cell size in proliferating cells (Magyar et al., 2005), but as opposed to SCL28 that solely acts on cell proliferation rate, the RBR-E2F pathway also regulates cell cycle exit, establishment of quiescence, and cellular differentiation (Borghi et al., 2010; Bouyer et al., 2018, Dewitte et al., 2003). The repressor-type MYB3Rs—MYB3R1, MYB3R3 and MYB3R5—when combined with RBR and E2FC form the DREAM complex that represses mitotic genes during or after developmental cessation of cell proliferation (Kobayashi et al., 2015). It appears that SCL28 acts independently of DREAM regulatory mechanisms, which explains why cell cycle exit and concomitant cell differentiation remain unaffected, but it controls cell cycle length specifically during proliferation.

In addition to cell size during proliferation, SCL28 also strongly affects the size of terminally differentiated cells. In the *scl28* mutant, we observed a reduced final cell size in various organs, whereas moderately higher *SCL28* expression led to its increase. Reduced cell size caused by increased cell division, as observed in *scl28*, has been repeatedly reported in Arabidopsis organ development. Examples include plants with inhibited *RBR* expression, those overexpressing *CYCD3*;*1*, or co-overexpressing *E2FA* and *DPA* (Dewitte et al., 2003; Borghi et al., 2010; Gutzat et al., 2011; De Veylder et al., 2002). In many of these cases, the cellular phenotype is due to prolonged cell proliferation and delayed cell differentiation, leading to additional cell division that is not balanced by cell size increase due to developmental decrease of cell expansion. On the contrary, we showed with kinematic analysis during leaf development that the increased number of smaller cells observed in *scl28* leaves is caused by accelerated cell proliferation, but not by prolonged cell proliferation. Therefore, it is puzzling how loss of *SCL28* affects final cell size that is largely determined by post-mitotic cell expansion occurring in the absence of SCL28.

Our observations are reminiscent of the phenomena reported in Drosophila wings, where the alteration of cell number is compensated by changes in cell size such that final organ size is essentially unchanged. Based on this and similar observations, it has been proposed that animal organ size may be controlled by ‘total mass checkpoint’ mechanisms, which operate at the level of whole organs (Potter and Xu, 2001), although the underlying molecular details remain unknown. Instead, the well-established mechanism of ‘compensation’ in plants shows postmitotic cell expansion that is enhanced by reduced cell proliferation (Hisanaga et al., 2015). A small number of larger cells observed in *SCL28*^OE^ leaves may be a prime example of compensation. One possible explanation for compensatory alteration of cell expansion is that, during cell proliferation, cells establish memory that is maintained during the elongation phase. An example of this can be seen in plants overexpressing *KRP2*, where final cell size may be influenced by the size of mitotic cells that are already altered during proliferation (Ferjani et al., 2007). We observed larger and smaller mitotic cells both in root meristem and young leaves of *SCL28*^OE^ and *slc28*, respectively, which may be maintained until terminal differentiation, thus affecting final cell size. A possible molecular mechanism to create such memory is cell size-dependent alteration in gene expression. It was shown in yeast that cell size changes due to ploidy levels are associated with altered gene expression related to functions of cell wall and extracellular matrix (Wu et al., 2010). Similarly, in Arabidopsis root, ploidy levels were correlated with expression of genes related to chromatin and cell expansion, such as ion transport and cell wall modifications (Bhosale et al., 2018). Therefore, it is plausible that SCL28 action in meristematic cells has long-lasting effects through cell expansion on final cell size in differentiated tissues. Another possible mechanism for cell size memory involves a monitoring system for cellular dimensions that shares components in proliferating and expanding cells. Supporting this idea, it has been suggested that cells expand to a target size by a mechanism requiring a cell size measurement device, from an approach combining quantitative analysis and mathematical modeling of variability in final cell size of Arabidopsis root (Pavelescu et al., 2018). Further careful studies are required to clarify how final cell size is affected by expression levels of SLC28 that is confined to proliferating cells.

In summary, we identified a novel hierarchical MYB3R-SCL28/AtSMOS1-SMR transcriptional regulatory pathway leading to cell cycle inhibition at G2/M that regulates cell size and number during organ development without dramatically altering organ size. This mechanism may help to adjust cell size to optimize cellular and tissue performance such as metabolic functions or mechanical properties.

## Methods

### Plant materials

*Arabidopsis thaliana* Columbia (Col) was used as the WT plant. All mutants and transgenic lines used in this study were in a Col background. The mutant alleles *scl28-1* (SALK_205284), *atsmos1-1* (SALK_111105C), *smr1-1* (SALK_033905), *smr2-1* (SALK_006098C), and *smr13-1* (SALK_053256C) were identified from SALK T-DNA collection, and used for phenotype analysis and generating multiple mutant combinations. Other mutants and transgenic lines, namely *myb3r1-1*, *myb3r3-1*, *myb3r4-1*, *myb3r5-1*, PCNA-RFP, and CYCB1;1-GFP, were described previously (Haga *et al*, 2007; Kobayashi et al., 2015; Yokoyama et al., 2016; Ubeda-Tomás et al., 2009). Sterilized seeds were germinated on one-half-strength Murashige and Skoog (1/2 MS) medium containing 2% (w/v) sucrose and 0.6% (w/v) agar. Plants were grown on 1/2 MS agar medium or soil under continuous light at 22°C. For root phenotype analysis, plants were gown on a vertical surface of 1/2 MS medium containing 1.0% (w/v) agar.

### Gene expression analysis

For comparing transcriptome of WT and *SCL28*^OE^, microarray analyses were performed using an ATH1 GeneChip^®^ (Affymetrix) as described previously (Kobayashi et al., 2015). For RNA-Seq analyses, cDNA libraries were constructed with the TruSeq RNA Library Preparation Kit v2 (Illumina, United States) and sequenced using the NextSeq500 sequencer (Illumina) as described previously (Okumura et al., 2021). RNA-Seq data for WT, *scl28*, and *atsmos1* can be accessed from the DDBJ database under accession number DRA012786. RNA extraction and qRT-PCR were performed as described previously (Okumura et al., 2021), using primers listed in Supplementary Table 1.

### Yeast two-hybrid assay

Yeast two-hybrid assays were performed as described by Soyano et al. (2003). *Saccharomyces cerevisiae* strain L40 was transformed with pBTM116- and pVP16-based plasmids carrying coding sequences encoding for SCL28 or AtSMOS1. A single colony was diluted with water and spotted on medium lacking His supplemented with 5 mM 3-AT and cultured at 30°C for two days.

### BiFC assay

For construction of plasmids used for BiFC assays, the entire coding regions of *SCL28* and *AtSMOS1* were amplified by PCR, and cloned into donor vectors pDONR201 and pDONR207 using BP Clonase II (Thermo Fisher), respectively. The cloned fragments were then transferred into the destination vectors pGWnY and pGWcY (Hino et al. 2011) using LR clonase II (Thermo Fisher) to generate C-terminal fusions of YFP fragments. Transient gene expression using Arabidopsis leaf mesophyll protoplasts was performed as described previously (Yoshida et al. 2014). The transfected protoplasts were incubated overnight in the dark at 22°C. YFP fluorescent was observed with fluorescent optics on a BX54 microscope (Olympus).

### LUC reporter assay

Preparation of protoplasts from Arabidopsis T87 suspension cultured cells and polyethylene glycol-mediated gene transfer were performed as described previously (Maeo et al. 2009). An empty vector containing 35S promoter (pJIT-60) and 35S:hRLUC plasmid expressing the humanized *Renilla* LUC (hRLUC) were used as a negative control and an internal control, respectively. Protoplasts (1.5 × 10^5^ for each transfection) were co-transfected with 15 *μ*g each of the *LUC* reporter and expression plasmids, then incubated at 22°C for 20 h before measuring LUC activities, which was performed using the Dual-Luciferase Reporter Assay system (Promega) and Luminoskan Ascent luminometer (Thermo Fisher). LUC activity was normalized according to hRLUC activity in each assay, and the relative ratio was determined.

### Plasmid construction

To construct GUS fusion reporters, the upstream region of *SCL28* (2.2 kb) was amplified by PCR and cloned into the pENTR/D-TOPO vector (Thermo Fisher), then transferred to pBGGUS (Haga et al., 2011) through Gateway LR reaction to create proSCL28::GUS. The PCR-based site-directed mutagenesis was performed to change all four MSA core motives, AACGG, in the *SCL28* promoter to ATTGG, resulting in generation of proSCL28ΔMSA::GUS.

To prepare the construct expressing SCL28 fused to GFP at its C-terminus (SCL28-GFP) under the control of the native promoter, the entire *SCL28* genomic region (5.3 kb) containing 2.2-kb promoter was amplified by PCR using genomic DNA from *Arabidopsis* (Col), and cloned into pENTR/D-TOPO. The resulting plasmid was then used for In-Fusion reaction (Takara Clontech) to insert PCR-amplified GFP fragments at the C-terminus of the *SCL28* coding sequence, to create entry plasmid containing the proSCL28:: SCL28-GFP fusion construct, which was then transferred to the binary vector pGWB501 (Nakagawa et al., 2007).

To prepare the AtSMOS1-GFP fusion construct driven by its own promoter, the entire *AtSMOS1* genomic region (4.4 kb) containing 1.2-kb promoter was amplified by PCR using genomic DNA from Arabidopsis (Col) and cloned into pDONR201 through Gateway BP reaction (Thermo Fisher). The resulting plasmid was then used for In-Fusion reaction (Takara Clontech) for inserting PCR-amplified GFP fragment at the C-terminus of *AtSMOS1*, to create entry plasmid containing proAtSMOS1::AtSMOS1-GFP fusion construct, which was then transferred to the binary vector pPZP211-GW (Kozgunova et al., 2016).

For construction of a LUC reporter plasmid, promoter region of *SMR2* (2.0 kb) was amplified by PCR and cloned into *HindIII-BamHI* interval of pUC-LUC (Ito et al., 2001) to obtain proSMR2::LUC. For construction of expression plasmids of AtSMOS1 and SCL28 (pro35S::AtSMOS1 and pro35S::SCL28), the entire coding regions were amplified by PCR using cDNA prepared from Arabidopsis T87 cells as a template. The resulting PCR fragments were cloned into pENTR/D-TOPO and then transferred to pJIT60 (Ito et al., 2001) at the site downstream of 35S promoter through the Gateway LR reaction.

To create proRPS5A::SCL28-GFP, the entire coding sequence of SCL28-GFP fusion was amplified by PCR using proSCL28::SCL28-GFP plasmid as a template and cloned into pENTR/D-TOPO. The resulting plasmid was then used for the Gateway LR reaction to transfer the SCL28-GFP fragment downstream of *RPS5A* promoter in pPZP221 (Haga et al., 2007). Primers used for plasmid construction are listed in Supplementary Table 1.

### Histological analysis

Excised plant organs were fixed in FAA solution (100% ethanol: formaldehyde: glacial acetic acid: water = 20: 19: 1: 1) under vacuum, stained with 1% Safranin O, dehydrated with 30%, 50%, 70%, and 100% ethanol series, and embedded in Technovit 7100 (Heraeus Kulzer). 2*μ*m-thickness sections were cut on RM2125RT microtome (Leica) equipped with a tungsten carbide disposable blade TC-65 (Leica). Sections were briefly stained in a 0.01%−0.5% (w/v) toluidine blue-O in 0.1% (w/v) Na_2_CO_3_ solution, then washed with 0.1% Na_2_CO_3_ solution. Sections were observed under BX63 microscope (Olympus) and images were acquired with cellSens Standard Software (Olympus).

### Chromatin immunoprecipitation assay

ChIP-Seq assays were performed on whole seedlings using anti-GFP antibody (Clontech 632592). Seedlings at 14 DAS (5 g) from proAtSMOS1::AtSMOS1-GFP and proSCL28::SCL28-GFP were crosslinked in 1% (v/v) formaldehyde at room temperature for 15 min. Crosslinking was then quenched with 0.125 M glycine for 5 min. The crosslinked seedlings were ground, and nuclei were isolated and lysed in nuclei lysis buffer (1% SDS, 50 mM Tris-HCl, 10 mM EDTA, pH 8.0). Crosslinked chromatin was sonicated using a water bath Bioruptor UCD-200 (Diagenode) (15sec on/15sec off pulses; 15 times). The complexes were immunoprecipitated with antibodies, overnight at 4°C with gentle shaking, and incubated for 1 h at 4°C with 40 *μ*L of Protein AG UltraLink Resin (Thermo Fisher). The beads were washed 2 × 5 min in ChIP Wash Buffer 1 (0.1% SDS, 1% Triton X-100, 20 mM Tris-HCl, 2 mM EDTA, 150 mM NaCl, pH 8.0), 2 × 5 min in ChIP Wash Buffer 2 (0.1% SDS, 1% Triton X-100, 20 mM Tris-HCl, 2 mM EDTA, 500 mM NaCl, pH 8.0), 2 × 5 min in ChIP Wash Buffer 3 (0.25 M LiCl, 1% NP-40, 1% sodium deoxycholate, 10 mM Tris-HCl, 1 mM EDTA, pH 8.0) and twice in TE (10 mM Tris-HCl, 1 mM EDTA, pH 8.0). ChIPed DNA was eluted by two 15-min incubations at 65°C with 250 μL elution buffer (1% SDS, 0.1 M NaHCO_3_). Chromatin was reverse-crosslinked by adding 20 μL of 5 M NaCl and incubated overnight at 65°C. Reverse-crosslinked DNA was subjected to RNase and proteinase K digestion and extracted with phenol-chloroform. DNA was ethanol precipitated in the presence of 20 μg of glycogen and resuspended in 50 μL of nuclease-free water (Ambion) in a DNA low-bind tube. We used 10 ng of IP or input DNA for ChIP-Seq library construction using NEBNext^®^ Ultra DNA Library Prep Kit for Illumina^®^ (New England Biolabs) according to manufacturer’s recommendations. For all libraries, twelve cycles of PCR were used. Library quality was assessed with Agilent 2100 Bioanalyzer (Agilent).

### Computational analysis of ChIP-Seq

Single-end sequencing of ChIP samples was performed using Illumina NextSeq 500 with a read length of 76 bp. Reads were quality controlled using FASTQC (http://www.bioinformatics.babraham.ac.uk/projects/fastqc/). Trimmomatic was used for quality trimming. Parameters for read quality filtering were set as follows: minimum length of 36 bp; mean Phred quality score greater than 30; and leading and trailing bases removal with base quality below 5. Reads were mapped onto the TAIR10 assembly using Bowtie (Langmead and Salzberg, 2012) with mismatch permission of 1 bp. To identify significantly enriched regions, we used MACS2 (Zhang et al., 2008). Parameters for peak detection were set as follows: number of duplicate reads at a location: 1; mfold of 5: 50; q-value cutoff: 0.05; extsize 200; and sharp peak. Visualization and analysis of genome-wide enrichment profiles were performed with IGB. Peak annotations such as proximity to genes and overlap of genomic features, including transposons and genes were performed using BEDTOOLS INTERSECT. NGSplot was used to profile enrichment at TSSs and along the gene (Shen et al., 2014). Spatial binding of the AtSMOS1 and SCL28 peaks were performed by position-wise comparison using a binning approach and plotted in hexplot. De novo motif analysis of both SCL28 and AtSMOS1 binding regions were screened using HOMER (Kim et al., 2020). ChIP-Seq data of SCL28 and AtSMOS1 can be accessed at Gene Expression Omnibus database under accession number GSE183209.

### Kinematic analysis of leaf growth

After plants were fixed in a 9:1 of ethanol and acetic acid solution, and cleared with Hoyer’s solution (a mixture of 100 g chloral hydrate, 10 g glycerol, 15 g gum arabic, and 25 ml water), we performed microscopic observations using 1st or 2nd leaves as described previously (Haga et al. 2007). After whole leaf images were captured, palisade cells at positions one-fourth and three-fourth from the tip of each leaf were observed with differential interference contrast (DIC) microscope (BX51, Olympus). The captured images were analyzed using ImageJ (ver.2.1.0; rsb.info.nih.gov/ij) and the average size of palisade cell, the number of cells in the uppermost layer of palisade tissue per leaf, and cell division rate were calculated according to methods described previously (De Veylder et al. 2001).

### Ploidy analysis

For ploidy analysis, nuclei were isolated from whole leaves from plants at 8−20 DAS, then stained using CyStain UV precise P kit (Sysmex). After filtration, samples were analyzed with CyFlow Ploidy Analyser (Sysmex) according to the manufacturer’s instructions. The population of nuclei in each peak was estimated as described previously (Kobayashi et al. 2015).

### Analysis of meristem cell number and cell size in roots

To visualize cell outlines, roots from plants at 5DAS were stained with 0.05 mg ml-1 propidium iodide and observed by confocal laser scanning microscopy (CLSM), using an inverted fluorescence microscope (Eclipse Ti2, Nikon) equipped with a confocal scanning unit (A1, Nikon). The resulting images were processed using ImageJ software to measure cell length in the root meristem. Root meristem size was measured by counting the number of cortical cells between the quiescent center and the first elongated cell.

### Cell cycle analysis by time-lapse imaging

Seedlings at 6 DAS carrying PCNA-GFP were transferred onto an MS medium in a glass-bottom dish, and observed by CLSM as described above. Time-lapse images were acquired every 30 min for 15 h. To avoid long-time imaging that possibly damages the samples, the duration of the G1/S and G2/M phases was measured separately.

### Other experimental procedures

ChIP-qPCR experiments (Kobayashi et al., 2015) and GUS staining (Haga et al., 2007) were performed as described previously. Primers used for ChIP-qPCR are listed in Supplementary Table 1.

## Data availability

Arabidopsis mutants and transgenic lines, as well as plasmids generated in this study are available from the corresponding author upon reasonable request.

## Acknowledgements

The authors thank Kyoko Kato, Akiko Nakanishi, Natsuko Ono, Akiko Yamamoto, Chikako Inoue, Nanako Ishibashi, Satoko Nasu, Yuiko Tachikawa, Asako Segawa, Ayami Furuta, Tomomi Shinagara and Ayumi Yamada for technical assistance. This study was funded by The Japan Society for the Promotion of Science KAKENHI (20H05408 and 17H03696 to M. Ito, 20H05911 and 20H05905 to K. Sugimoto), and bythe Institut Universitaire de France (to M. Benhamed). Y. Huang was supported by China Scholar Council fellowships (201806690005).

## Author contributions

M. Ito. conceived the study; C.R., M.B., Z.M., K.S., T.Y., and M.I. designed the experiments; M. Imamura., K.M., T.I., C.B., M.G., D.L., Y.H., Toshiya Suzuki, K.Y., H.T., and Y.N. carried out the experiments; J.A., Takamasa Suzuki analyzed the data; M.I., L.B., Z.M., and M.B. wrote the paper

## Additional information

**Supplementary information** The online version contains supplementary material.

