## Supplementary Figures and Tables for "A hierarchical transcriptional network activates specific CDK inhibitors that regulate G2 to control cell size and number in Arabidopsis"

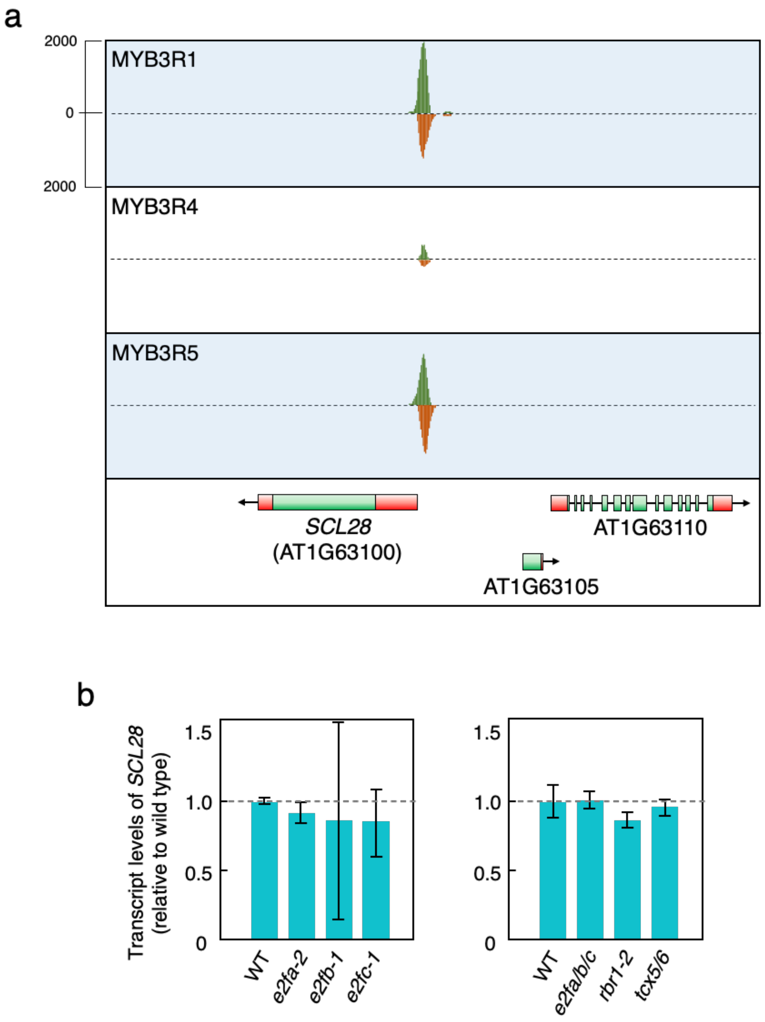


**Supplementary Figure 1**

**Profiles of DAP-seq around *SCL28* locus.**

**(a)** The DAP-seq profiles for MYB3R1, MYB3R4, and MYB3R5 were obtained by the Plant Cistrome Database at http://neomorph.salk.edu/dap_web/pages/index.php (O'Malley et al., 2016). Peaks shown in green and orange correspond to DAP-seq reads of forward and reverse orientations, respectively.

**(b)** Transcript levels of *SCL28* in the mutants lacking genes for DREAM components. qRT-PCR was performed using whole seedlings of WT, *e2fa-2*, *e2b-1*, and *e2fc-1* (left) and those of WT, *e2fa/b/c*, *rbr1-2*, and *tcx5/6* (right). There was no significant difference (Student's t-test, $P < 0.05$) between WT and any of mutants examined. All mutants and mutant combinations have been described previously (Nowack et al., 2012; Wang et al., 2014; Lang et al., 2121).

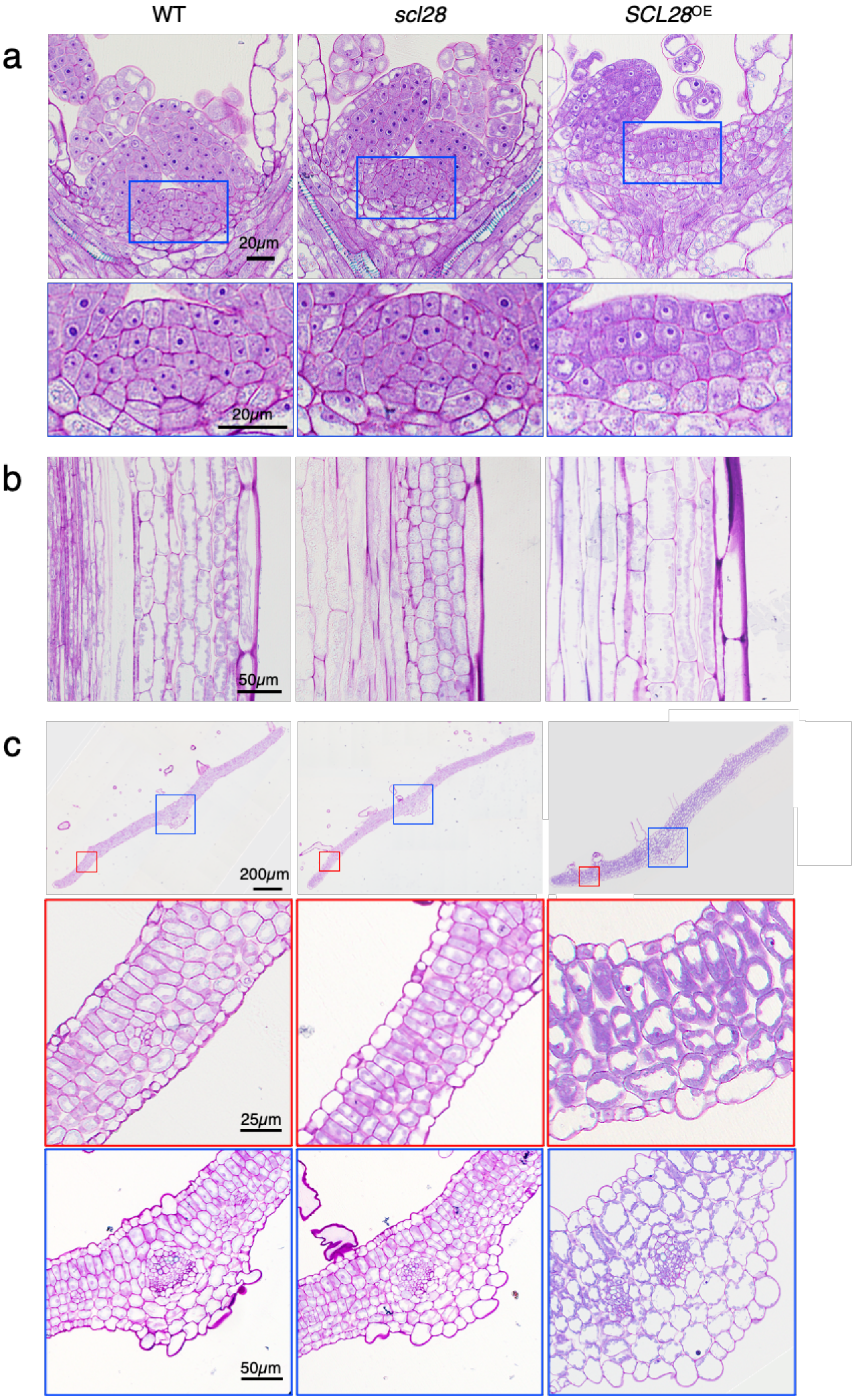

WT
scl28
SCL28OE
a
20μm
20μm
b
50μm
c
200μm
25μm
50μm

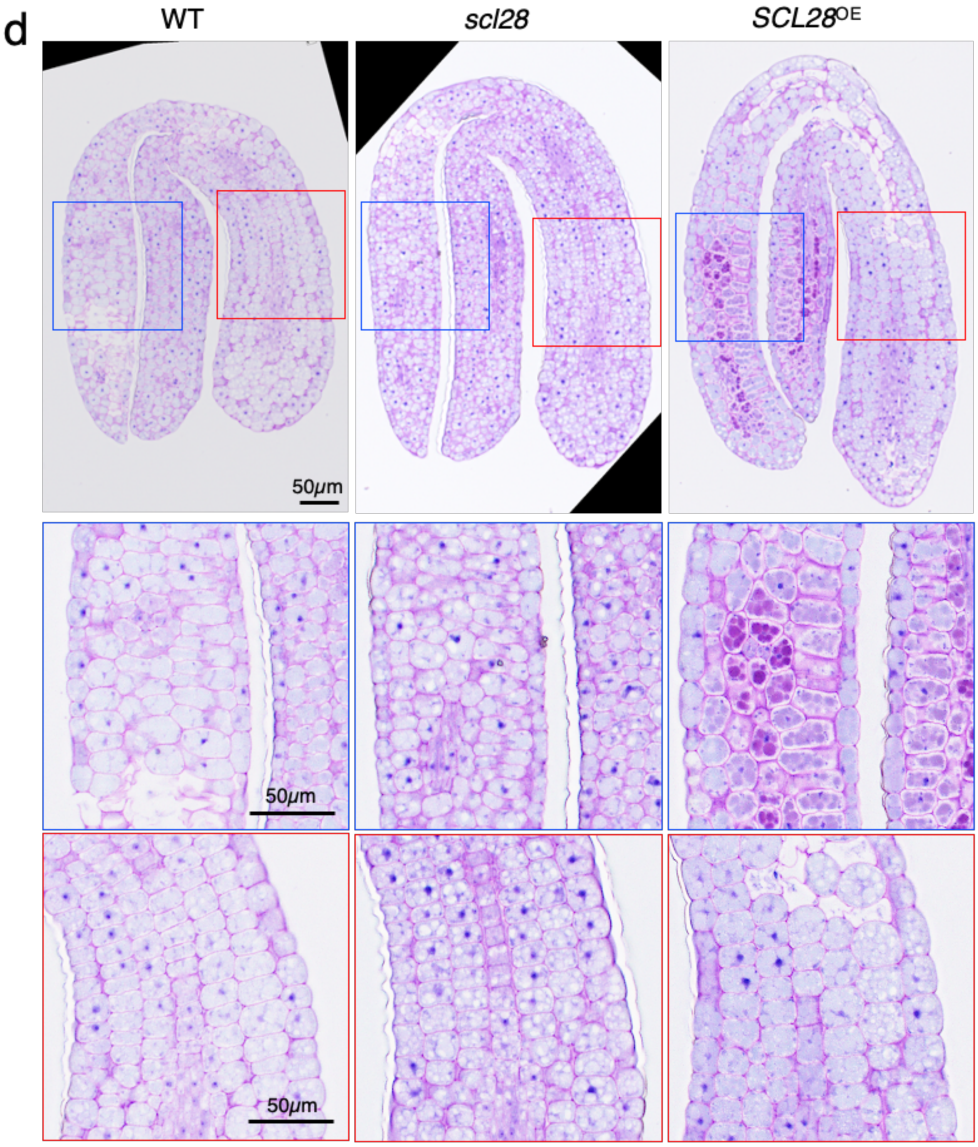


**Supplementary Figure 2**

**Cell size in various organs is oppositely affected in *scl28* and *SCL28*$^{OE}$ plants.**

**(a)** Longitudinal sections of shoot apical meristem and leaf primordia. The area shown in blue rectangles are magnified, and shown in lower panels.

**(b)** Vertical sections of inflorescence stems.

**(c)** Transverse sections of leaves. Areas surrounded by red and blue rectangles are magnified and shown in lower panels.

**(d)** Longitudinal sections of mature embryos. Regions surrounded by red and blue rectangles are magnified and shown in lower panels.

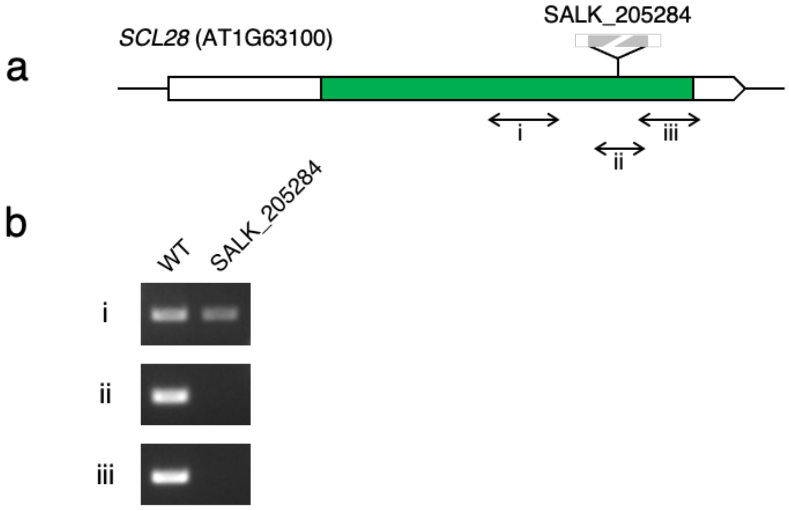


**Supplementary Figure 3**

**Confirmation of T-DNA insertion mutation (SALK_205284) to be a null allele of *SCL28*.**

**(a)** Position of T-DNA insertion in the *SCL28* gene. Exons are shown by boxes, and introns by lines between boxes, where green and white boxes indicate coding and non-coding regions, respectively. Amplified regions by semi-quantitative PCR analysis are shown by double-headed arrows.

**(b)** Semi-quantitative PCR was performed using primer pairs sandwiching the regions (i, ii, and iii) indicated in (**a**).

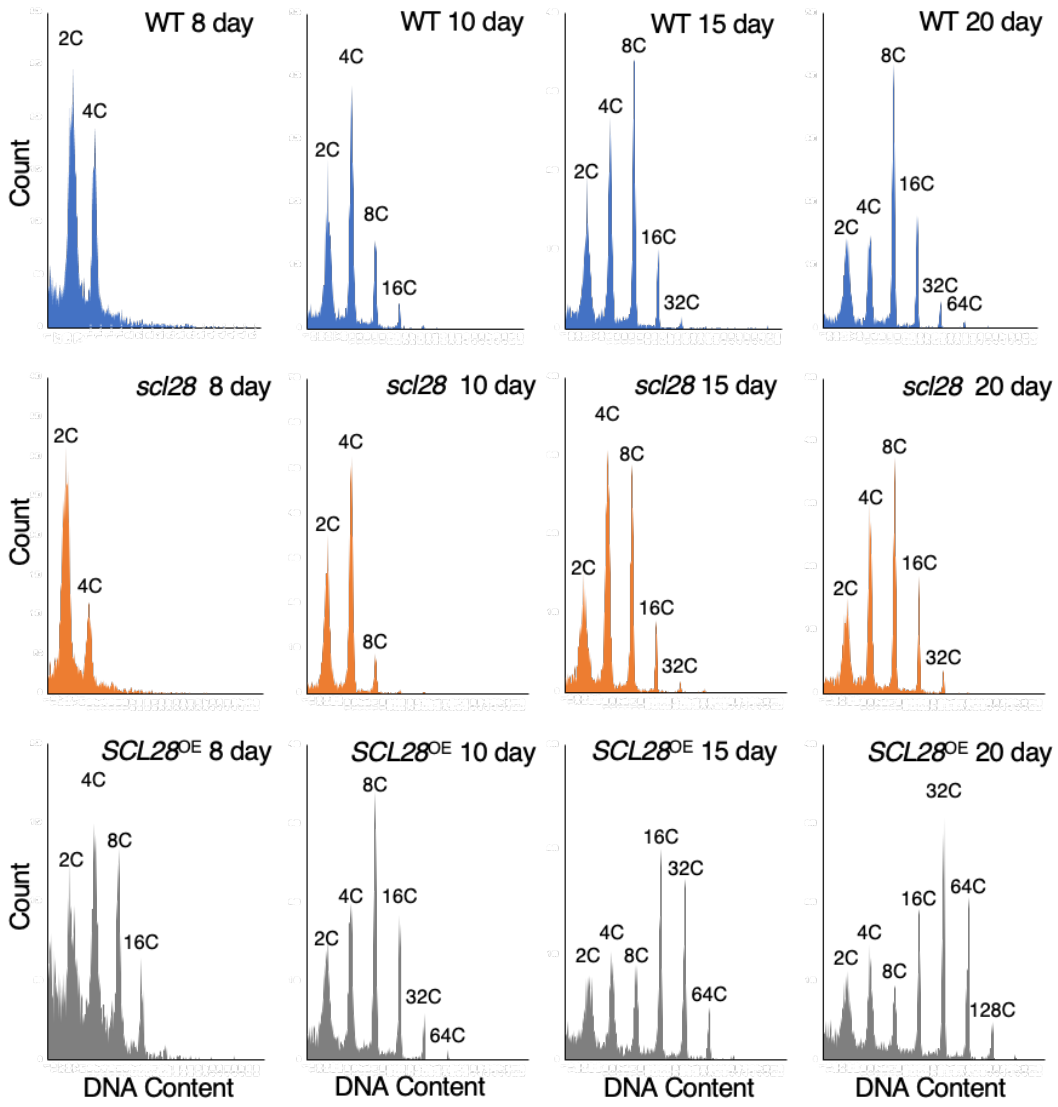


**Supplementary Figure 4**

**Ploidy analysis of WT, *scl28*, and *SCL28*$^{OE}$ plants.**

Representative profiles of ploidy distribution in first leaf pairs from WT, *scl28*, and *SCL28*$^{OE}$ plants grown for indicated period after sowing.

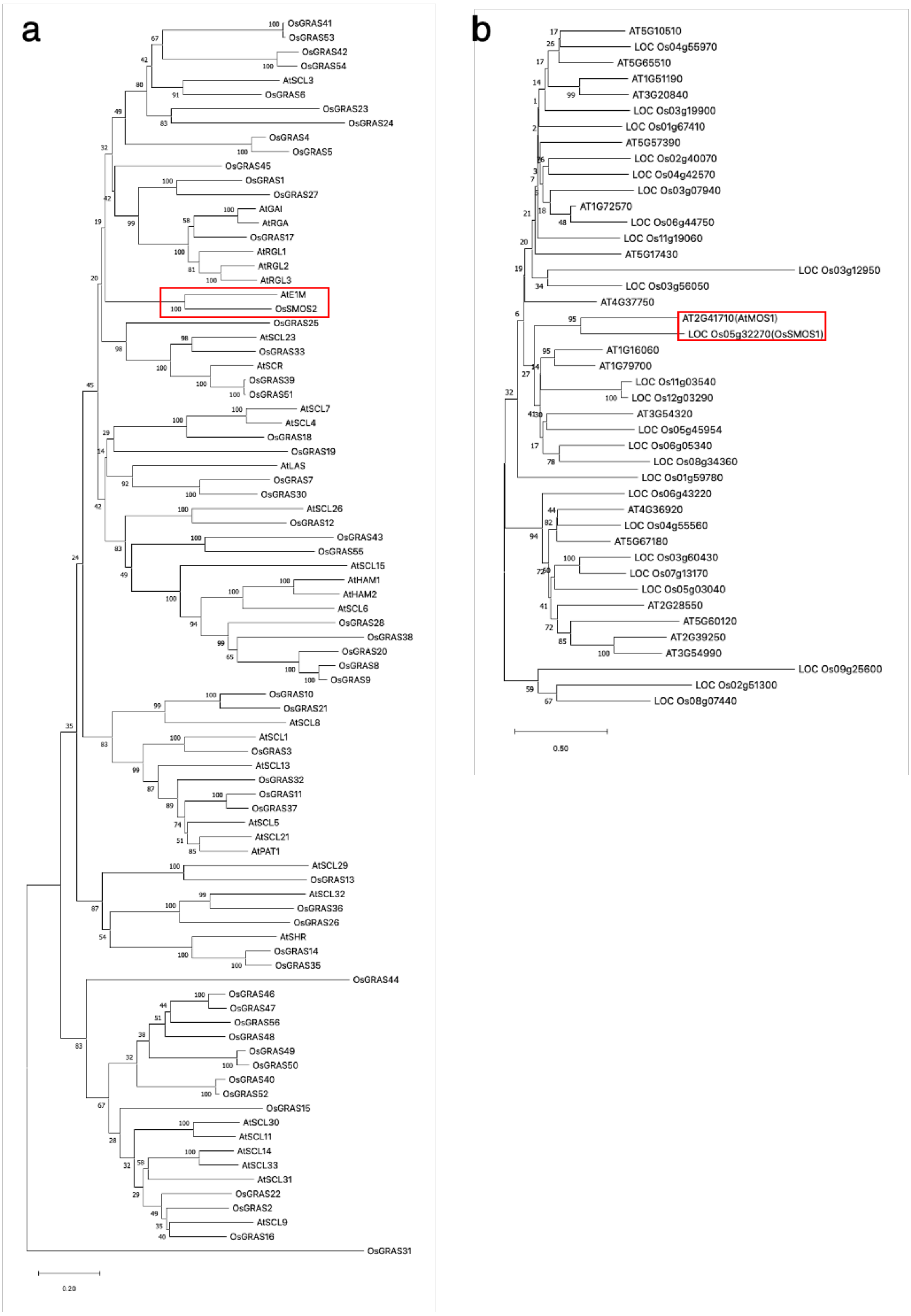


**Supplementary Figure 5**

**Phylogenetic analysis of SCL28 and AtSMOS1**

**(a)** Phylogenetic analysis of GRAS family proteins from rice and *Arabidopsis*, showing that SCL28 is orthologous to rice SMOS2. Amino acid sequences of all GRAS proteins in rice and *Arabidopsis* were obtained in Phytozome at https://phytozome.jgi.doe.gov/pz/portal.html (Goodstein et al., 2012). Amino acid sequence alignment within GRAS domain was generated by the MUSCLE program (Edgar 2004), and used for creating phylogenetic trees using MEGAX (Kumar et al. 2018) based on neighbor-joining method (Saitou and Nei, 1987).

**(b)** Phylogenetic analysis of AP2-type transcription factors from rice and Arabidopsis, showing that At2g41710 (AtSMOS1) is orthologous to rice SMOS1. Amino acid sequences of Arabidopsis proteins categorized as AP2 subfamily (Dietz et al., 2010) and corresponding rice proteins (Sharoni et al., 2011) were obtained and analyzed as in (**a**).

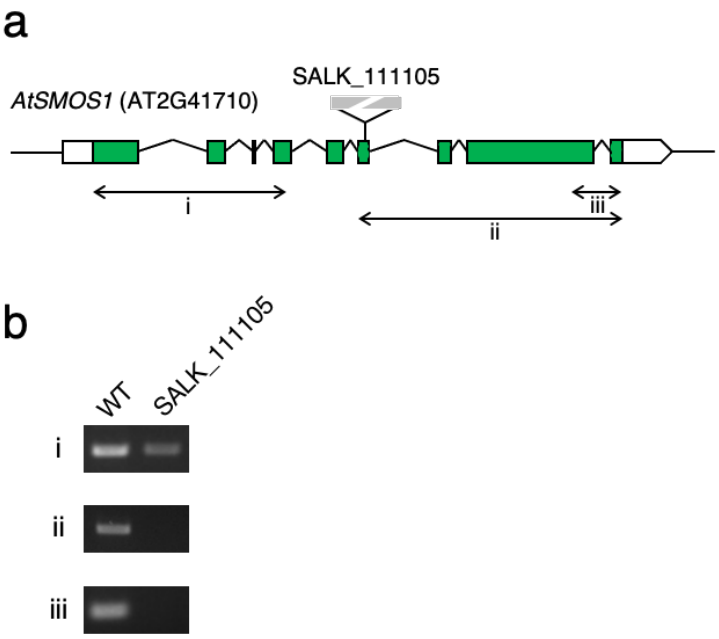


**Supplementary Figure 6**

**Confirmation of T-DNA insertion mutation (SALK_111105) to be a null allele of *AtSMOS1*.**

(**a**) Structure of *AtSMOS1* gene (At2g41710) and the position of T-DNA insertion. Exons are shown by boxes and introns by lines between boxes, where green and white boxes indicate coding and non-coding regions, respectively. Amplified regions by semi-quantitative PCR analysis are shown by double-headed arrows.

(**b**) Semi-quantitative PCR was performed using primer pairs sandwiching the regions (i, ii, and iii) indicated in (**a**).

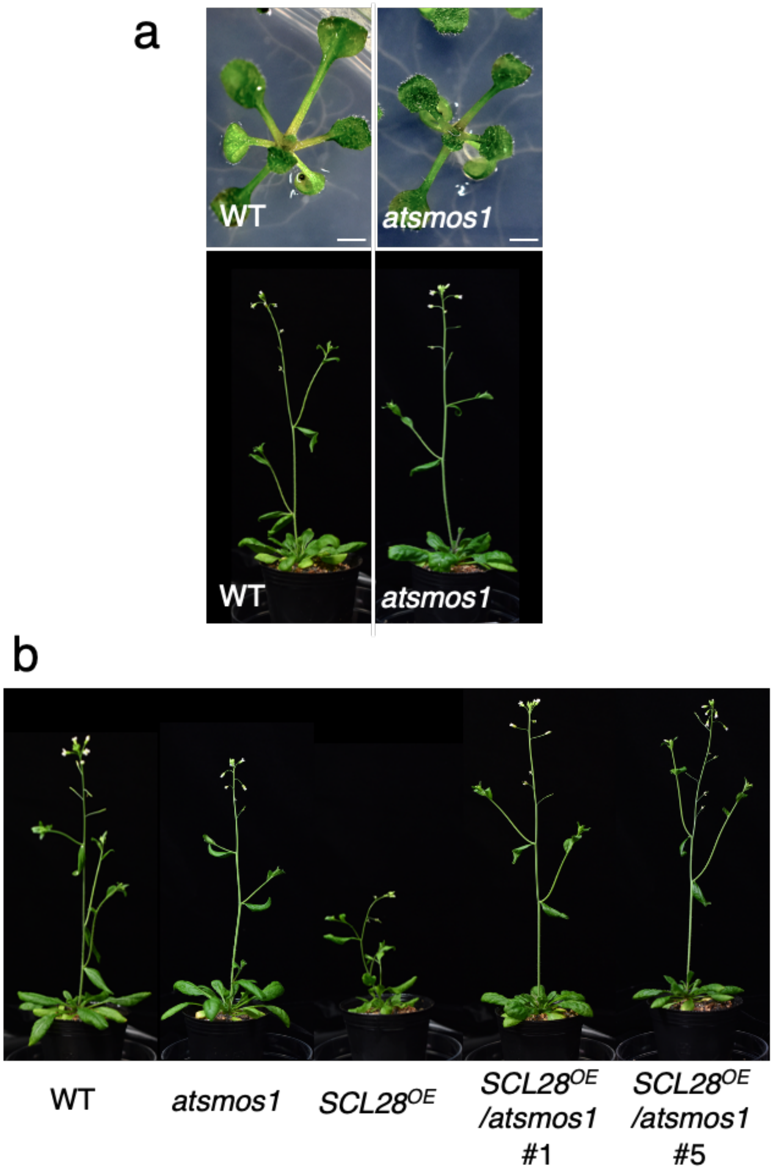


**Supplementary Figure 7**

***atsmos1* mutation does not significantly affect the plant growth, but completely suppress the growth inhibition caused by *SCL28* overexpression**

**(a)** Comparison of whole plant appearance between WT and *atsmos1*. Plants grown for 12 days on agar medium (upper) and those grown for four weeks on soil (lower) were photographed. Scale bars indicate 1 mm in upper panels.

**(b)** Comparison of whole plant appearance between WT, *atsmos1*, and *SCL28*$^{OE}$ plants and those carrying *atsmos1* and *SCL28*$^{OE}$ in combination (*SCL28*$^{OE}$/ *atsmos1*). Plants with indicated genotypes were photographed four weeks after sowing. Two independent lines of *SCL28*$^{OE}$/ *atsmos1* were analyzed.

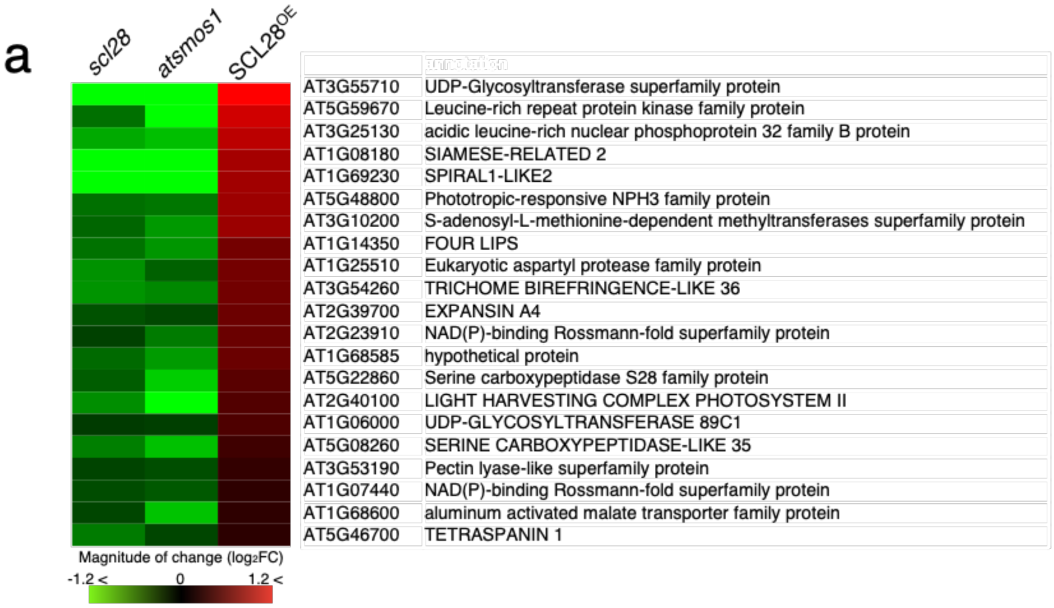


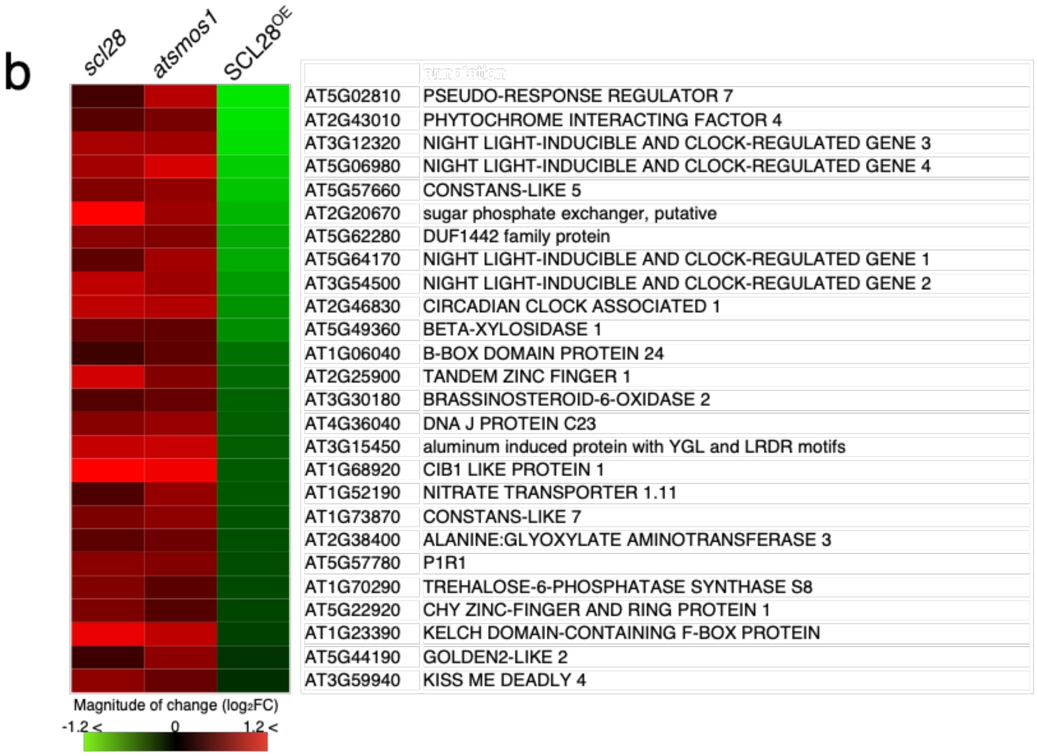


**Supplementary Figure 8**

**Heatmap representation of candidate genes regulated by SCL28-AtSMOS1 complex.**

**(a)** Heatmap showing expression changes of 21 genes downregulated in both *scl28* and *atsmos1* and upregulated in *SCL28*$^{OE}$.

**(b)** Heatmap showing expression changes of 26 genes upregulated in both *scl28* and *atsmos1* and downregulated in *SCL28*$^{OE}$.

Fold change level (log2) of each gene was calculated by comparing expression levels in *scl28*, *atsmos1*, or *SCL28*$^{OE}$ plants with that in WT plants.

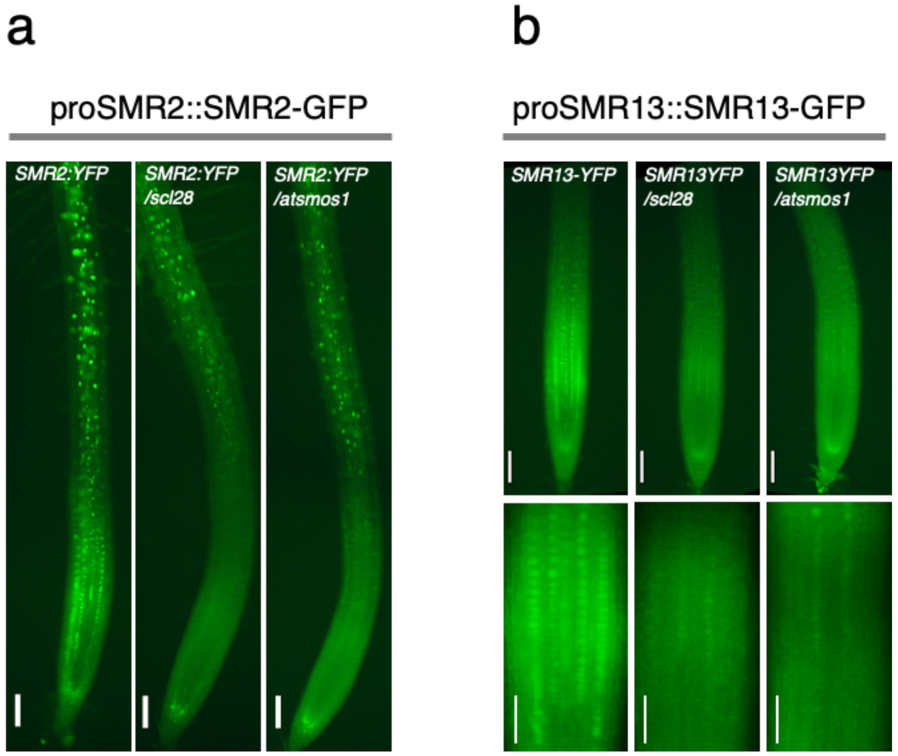


**Supplementary Figure 9**

**Downregulation of SMR2-GFP and SMR13-GFP in *scl28* and *atsmos1* mutants.**

**(a)** Fluorescent images of primary roots from plants at 7 DAS carrying proSMR2::SMR2-GFP under WT, *scl28* or *atsmos1* background. Scale bars indicate 100 $\mu$m

**(b)** Fluorescent images of primary roots from plants at 7 DAS carrying proSMR13::SMR13-GFP under wild type, *scl28* or *atsmos1* background. Magnified views of meristematic regions are shown in lower panels. Scale bars indicate 100 $\mu$m (upper) and 50 $\mu$m (lower).

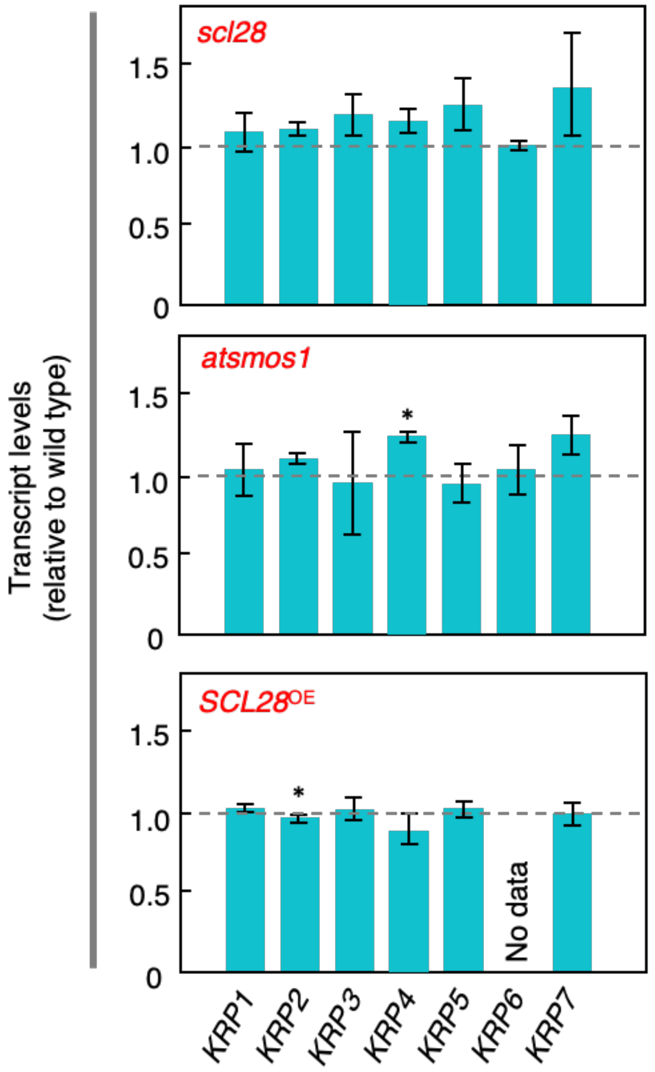


**Supplementary Figure 10**

**Transcript levels of *KRP* genes are not largely affected by *SCL28* and *AtSMOS1*.**

Expression changes of seven *KRP* genes in *scl28*, *atsmos1*, and *SCL28*$^{OE}$ plants. Relative expression levels of each *KRP* gene compared with those in WT are calculated, and shown as mean ± SD ($n = 3$). Statistical significance compared with WT was determined using Student's t-test. *$P < 0.05$

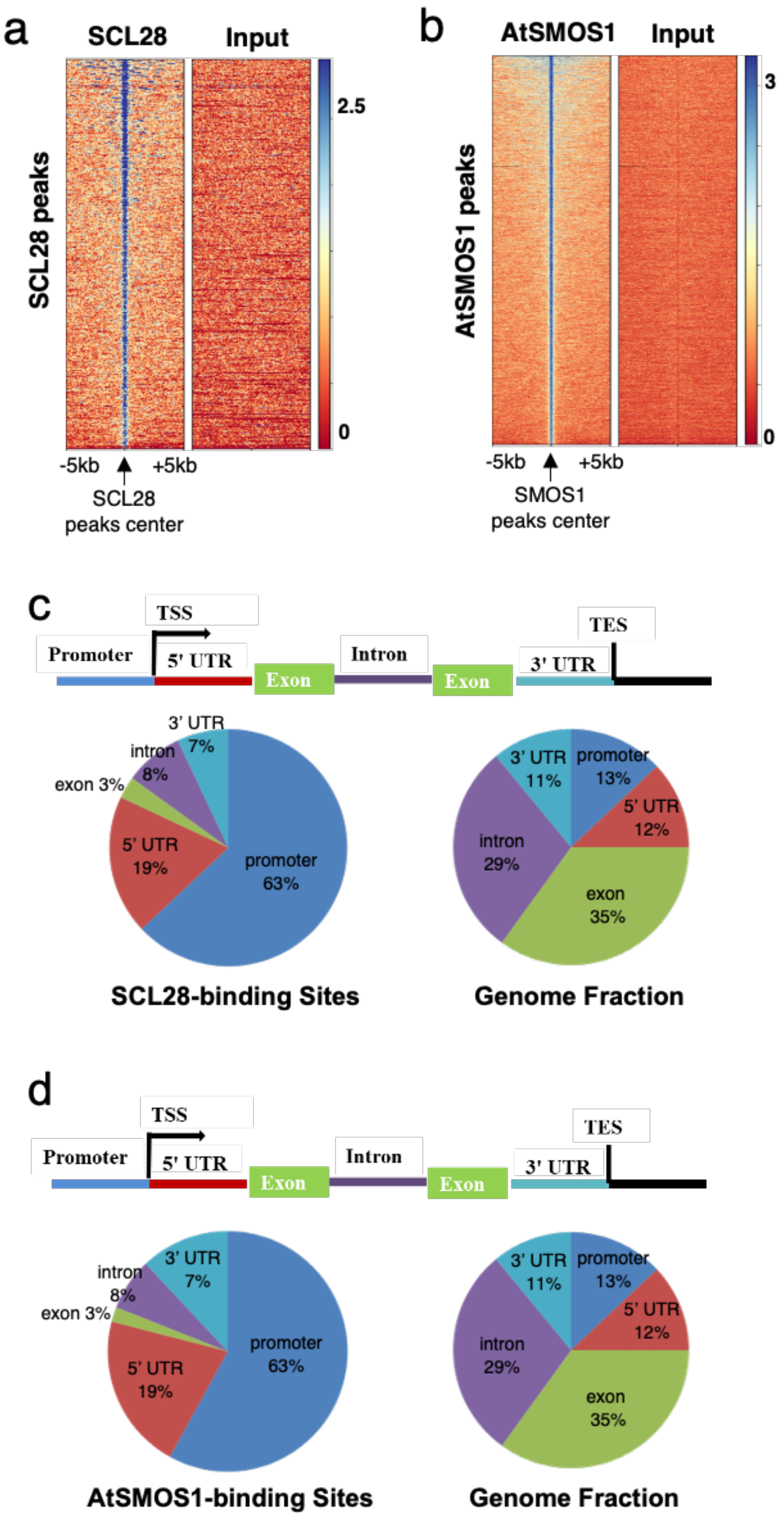


**Supplementary Figure 11**

**SCL28 and AtSMOS1 preferentially bind to promoter regions.**

**(a)** Comparison between SCL28 and input of tag density in the ±5 kb region around the SCL28 peaks showing

ChIP-seq peaks of SCL28 were successfully detected.

**(b)** Comparison between AtSMOS1 and input of tag density in the ±5 kb region around the AtSMOS1 peaks showing ChIP-seq peaks of AtSMOS1 were successfully detected.

**(c)** Pie chart representation of the distribution of SCL28 peaks identified by ChIP-seq in different genomic regions. The definition of each region is described above the pie chart.

**(d)** Pie chart representation of the distribution of AtSMOS1 peaks identified by ChIP-seq in different genomic regions. The definition of each region is described above the pie chart.

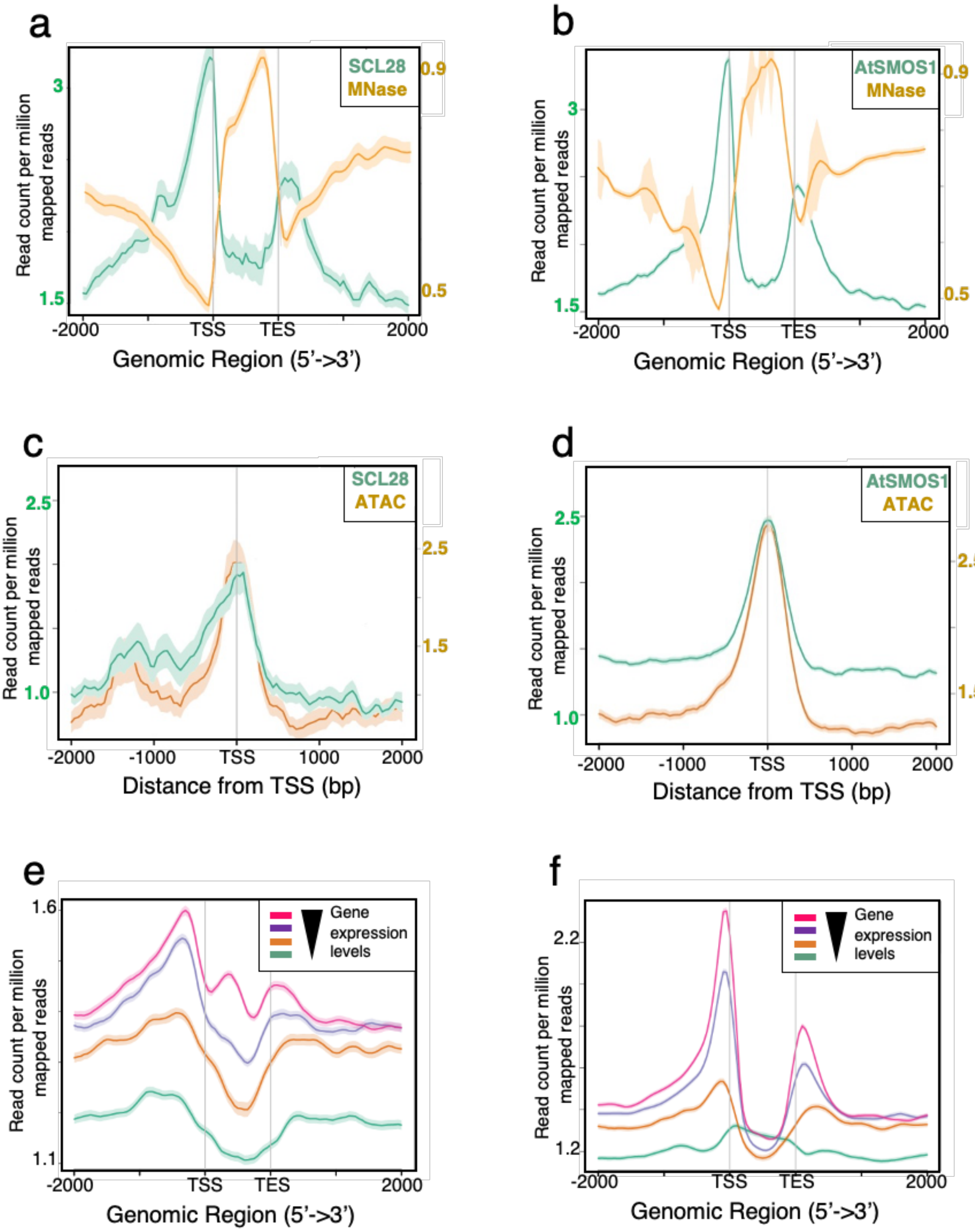


**Supplementary Figure 12**

**SCL28 and AtSMOS1 bind to nucleosome-free and highly accessible chromatin regions with their enrichment levels associated with mRNA levels**

**(a)** SCL28 binds nucleosome-free regions of transcribed genes. Mean profile of SCL28 ChIP-seq and MNase-seq reads density with respect to a gene model from TSS to TES. Normalization of coverage using spline algorithm was performed over the genes and flanking 2 kb region.

**(b)** AtSMOS1 binds nucleosome-free regions of transcribed genes. Data from AtSMOS1 ChIP-seq and MNase-seq were analyzed and shown as in (**a**).

**(c)** SCL28 binds to chromatin accessible sites. Profiles of SCL28 ChIP-seq and ATAC-seq reads density were merged over TSS and flanking 2 kb region.

**(d)** AtSMOS1 binds to chromatin accessible sites. Profiles of AtSMOS1 ChIP-seq and ATAC-seq reads density were merged over TSS and flanking 2 kb region.

**(e)** Highly-expressed genes show higher enrichment for SCL28 binding. Average enrichment profile of SCL28 is correlated with gene expression variations. Gene expression is categorized from low to high expression. Mean-normalized ChIP-Seq densities of equal bins were plotted along the gene and 2-kb region flanking the TSS or TES.

**(f)** Highly-expressed genes show higher enrichment for AtSMOS1 binding. ChIP-Seq data of AtSMOS1 were analyzed as in (**e**).

| | GO biological process | Arabidopsis thaliana REFLIST | # | Expected | over/ under | Fold Enrichment | raw P-value | FDR |
|---|---|---|---|---|---|---|---|---|
| ● | negative regulation of mitotic nuclear division (GO:0045839) | 25 | 5 | 0.17 | + | 28.76 | 1.85E-06 | 5.50E-03 |
| ● | negative regulation of nuclear division (GO:0051784) | 30 | 5 | 0.21 | + | 23.96 | 4.09E-06 | 4.87E-03 |
| ● | regulation of mitotic nuclear division (GO:0007088) | 36 | 5 | 0.25 | + | 19.97 | 9.14E-06 | 9.06E-03 |
| ● | regulation of DNA endoreduplication (GO:0032875) | 37 | 5 | 0.26 | + | 19.43 | 1.03E-05 | 8.76E-03 |
| | water transport (GO:0006833) | 30 | 4 | 0.21 | + | 19.17 | 8.70E-05 | 3.04E-02 |
| | fluid transport (GO:0042044) | 30 | 4 | 0.21 | + | 19.17 | 8.70E-05 | 2.87E-02 |
| ● | regulation of DNA-dependent DNA replication (GO:0090329) | 47 | 5 | 0.33 | + | 15.3 | 2.98E-05 | 2.22E-02 |
| ● | negative regulation of mitotic cell cycle (GO:0045930) | 48 | 5 | 0.33 | + | 14.98 | 3.27E-05 | 2.16E-02 |
| ● | regulation of nuclear division (GO:0051783) | 50 | 5 | 0.35 | + | 14.38 | 3.92E-05 | 2.33E-02 |
| | negative regulation of organelle organization (GO:0010639) | 50 | 5 | 0.35 | + | 14.38 | 3.92E-05 | 2.12E-02 |
| ● | regulation of DNA replication (GO:0006275) | 55 | 5 | 0.38 | + | 13.07 | 5.99E-05 | 2.37E-02 |
| ● | negative regulation of cell cycle process (GO:0010948) | 68 | 5 | 0.47 | + | 10.57 | 1.53E-04 | 4.14E-02 |
| ● | regulation of mitotic cell cycle (GO:0007346) | 86 | 6 | 0.6 | + | 10.03 | 4.37E-05 | 2.17E-02 |
| | response to water (GO:0009415) | 376 | 11 | 2.62 | + | 4.21 | 8.55E-05 | 3.18E-02 |
| | response to acid chemical (GO:0001101) | 408 | 11 | 2.84 | + | 3.88 | 1.71E-04 | 4.43E-02 |
| | response to oxygen-containing compound (GO:1901700) | 1431 | 28 | 9.95 | + | 2.81 | 9.52E-07 | 5.67E-03 |
| | oxoacid metabolic process (GO:0043436) | 976 | 18 | 6.79 | + | 2.65 | 1.96E-04 | 4.66E-02 |
| | cellular response to chemical stimulus (GO:0070887) | 1117 | 20 | 7.77 | + | 2.57 | 1.26E-04 | 3.96E-02 |
| | response to external stimulus (GO:0009605) | 1527 | 24 | 10.62 | + | 2.26 | 2.02E-04 | 4.62E-02 |
| | response to chemical (GO:0042221) | 2530 | 39 | 17.6 | + | 2.22 | 2.26E-06 | 4.49E-03 |
| | regulation of cellular macromolecule biosynthetic process (GO:2000112) | 2374 | 33 | 16.51 | + | 2 | 1.42E-04 | 4.21E-02 |
| | regulation of macromolecule biosynthetic process (GO:0010556) | 2387 | 33 | 16.6 | + | 1.99 | 1.51E-04 | 4.28E-02 |
| | regulation of cellular biosynthetic process (GO:0031326) | 2488 | 34 | 17.3 | + | 1.96 | 1.94E-04 | 4.82E-02 |
| | regulation of biosynthetic process (GO:0009889) | 2525 | 34 | 17.56 | + | 1.94 | 2.17E-04 | 4.78E-02 |
| | response to stress (GO:0006950) | 3187 | 42 | 22.17 | + | 1.89 | 5.70E-05 | 2.42E-02 |
| | response to stimulus (GO:0050896) | 5474 | 66 | 38.07 | + | 1.73 | 2.73E-06 | 4.06E-03 |
| | cellular process (GO:0009987) | 12044 | 112 | 83.77 | + | 1.34 | 5.16E-05 | 2.36E-02 |

● Cell cycle-related GO terms

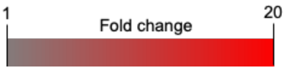


**Supplementary Figure 13**

**Gene ontology enrichment analysis of common targets of SCL28 and AtSMOS1.**

GO enrichment analysis was performed using PANTHER overrepresentation test with the "GO biological processes complete" dataset. Overrepresented GO terms related to cell cycle are shown by red dots on the left. Blue backgrounds show the number of common target genes belonging to the corresponding cell cycle-related GO categories, all of which include five common *SMR* genes (*SMR2*, *SMR4*, *SMR6*, *SMR8*, and *SMR9*).

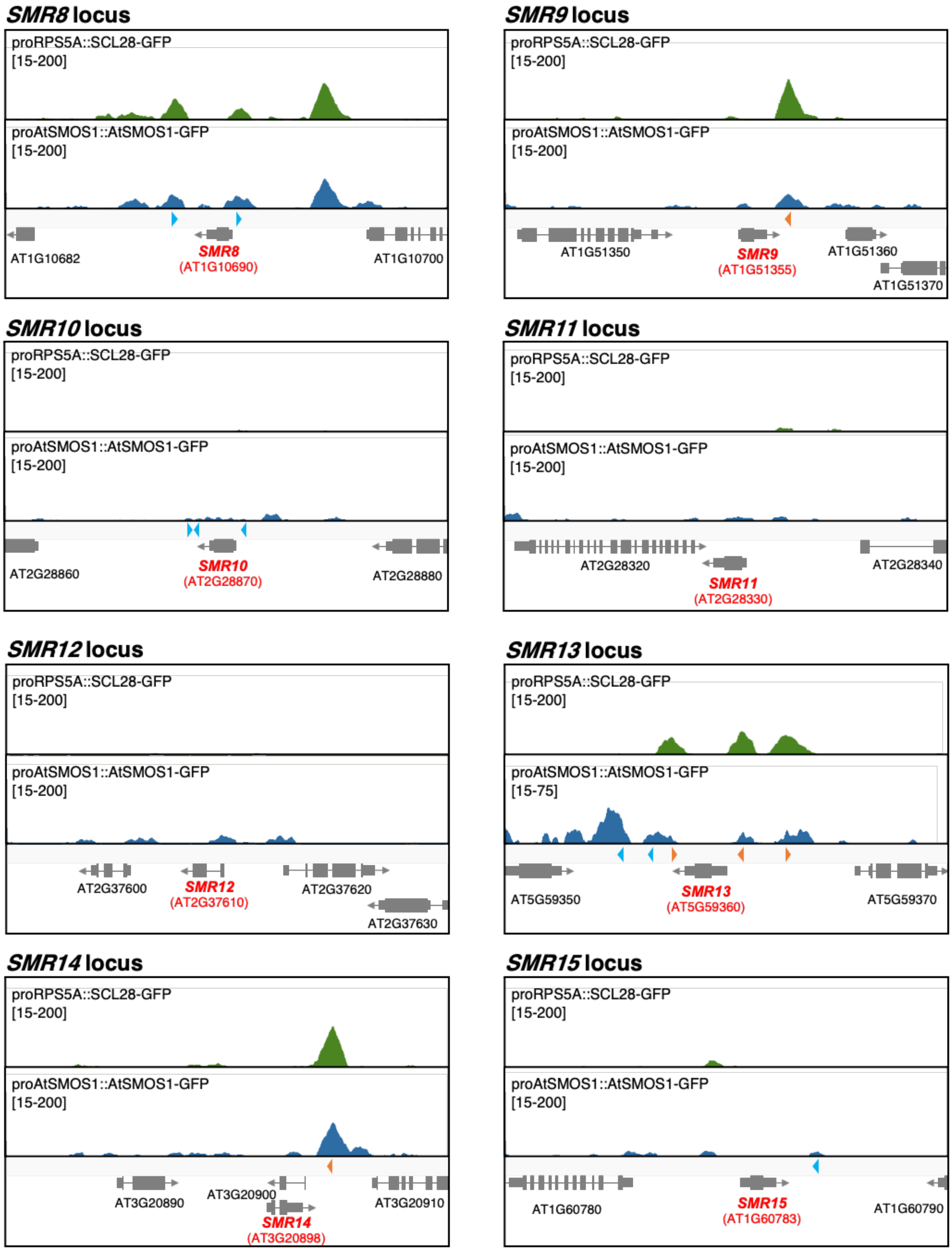
SMR8 locus
proRPS5A::SCL28-GFP
[15-200]
proAtSMOS1::AtSMOS1-GFP
[15-200]
AT1G10682
SMR8
(AT1G10690)
AT1G10700
SMR9 locus
proRPS5A::SCL28-GFP
[15-200]
proAtSMOS1::AtSMOS1-GFP
[15-200]
AT1G51350
SMR9
(AT1G51355)
AT1G51360
AT1G51370
SMR10 locus
proRPS5A::SCL28-GFP
[15-200]
proAtSMOS1::AtSMOS1-GFP
[15-200]
AT2G28860
SMR10
(AT2G28870)
AT2G28880
SMR11 locus
proRPS5A::SCL28-GFP
[15-200]
proAtSMOS1::AtSMOS1-GFP
[15-200]
AT2G28320
SMR11
(AT2G28330)
AT2G28340
SMR12 locus
proRPS5A::SCL28-GFP
[15-200]
proAtSMOS1::AtSMOS1-GFP
[15-200]
AT2G37600
SMR12
(AT2G37610)
AT2G37620
AT2G37630
SMR13 locus
proRPS5A::SCL28-GFP
[15-200]
proAtSMOS1::AtSMOS1-GFP
[15-75]
AT5G59350
SMR13
(AT5G59360)
AT5G59370
SMR14 locus
proRPS5A::SCL28-GFP
[15-200]
proAtSMOS1::AtSMOS1-GFP
[15-200]
AT3G20890
AT3G20900
SMR14
(AT3G20898)
AT3G20910
SMR15 locus
proRPS5A::SCL28-GFP
[15-200]
proAtSMOS1::AtSMOS1-GFP
[15-200]
AT1G60780
SMR15
(AT1G60783)
AT1G60790

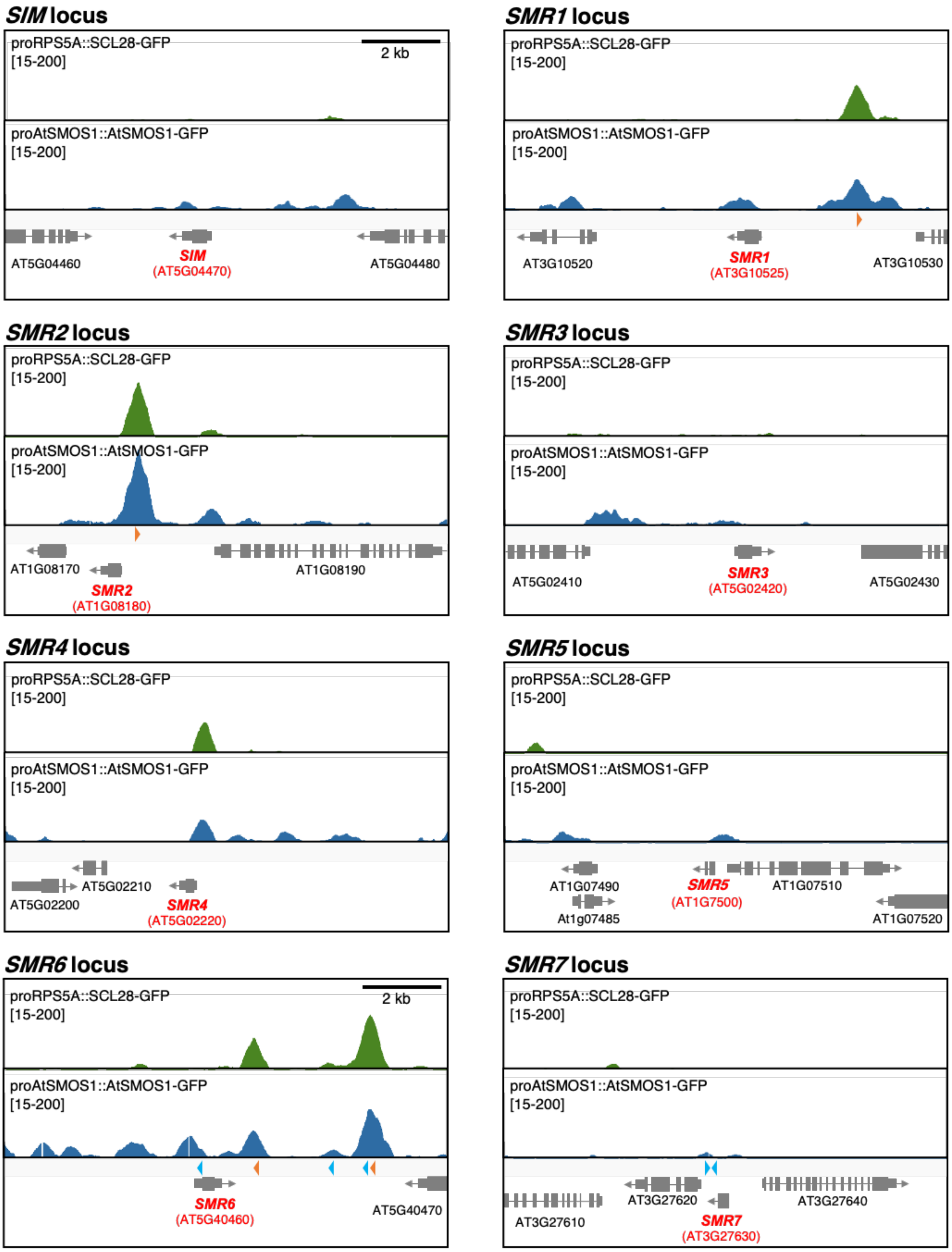
SIM locus
proRPS5A::SCL28-GFP
[15-200]
2 kb
proAtSMOS1::AtSMOS1-GFP
[15-200]
AT5G04460
SIM
(AT5G04470)
AT5G04480
SMR1 locus
proRPS5A::SCL28-GFP
[15-200]
proAtSMOS1::AtSMOS1-GFP
[15-200]
AT3G10520
SMR1
(AT3G10525)
AT3G10530
SMR2 locus
proRPS5A::SCL28-GFP
[15-200]
proAtSMOS1::AtSMOS1-GFP
[15-200]
AT1G08170
SMR2
(AT1G08180)
AT1G08190
SMR3 locus
proRPS5A::SCL28-GFP
[15-200]
proAtSMOS1::AtSMOS1-GFP
[15-200]
AT5G02410
SMR3
(AT5G02420)
AT5G02430
SMR4 locus
proRPS5A::SCL28-GFP
[15-200]
proAtSMOS1::AtSMOS1-GFP
[15-200]
AT5G02210
AT5G02200
SMR4
(AT5G02220)
SMR5 locus
proRPS5A::SCL28-GFP
[15-200]
proAtSMOS1::AtSMOS1-GFP
[15-200]
AT1G07490
At1g07485
SMR5
(AT1G7500)
AT1G07510
AT1G07520
SMR6 locus
proRPS5A::SCL28-GFP
[15-200]
2 kb
proAtSMOS1::AtSMOS1-GFP
[15-200]
SMR6
(AT5G40460)
AT5G40470
SMR7 locus
proRPS5A::SCL28-GFP
[15-200]
proAtSMOS1::AtSMOS1-GFP
[15-200]
AT3G27610
AT3G27620
SMR7
(AT3G27630)
AT3G27640

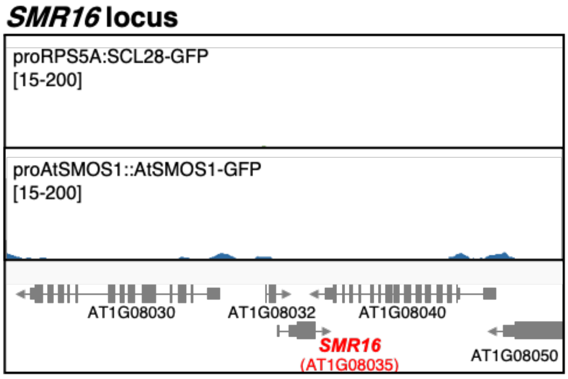


**Supplementary Figure 14**

**ChIP-seq profile of SCL28 and AtSMOS1 around the *SMR* genes.**

Genome-wide binding of SCL28 and AtSMOS1 was analyzed by ChIP-seq using proRPS5A::SCL28-GFP and proAtSMOS1::AtSMOS1-GFP plants. The ChIP-seq profiles of SCL28 and AtSMOS1 are shown for all 17 genes of the SMR family in Arabidopsis. Orange arrowheads indicate DNA motifs perfectly matching C(a/t)T(a/t)GGATNC(c/t)(a/t) identified as an enriched motif in the SCL28/AtSMOS1 common targets, whereas blue arrowheads indicate motifs matching the enriched motif with one base mismatch.

a

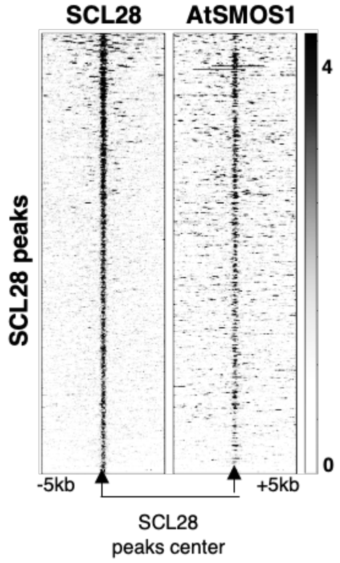


b

| Motif | *P*-value | log *P*-value |
| --- | --- | --- |
| 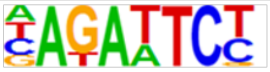 | 1e-31 | -7.142e+01 |
| 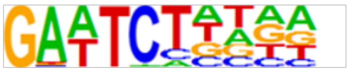 | 1e-27 | -6.327e+01 |
| 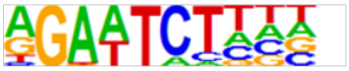 | 1e-26 | -6.107e+01 |

c

| Motif | *P*-value | log *P*-value |
| --- | --- | --- |
| 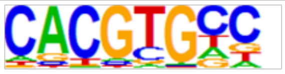 | 1e-241 | -5.553e+02 |

**Supplementary Figure 15**

**Supporting evidence for binding of SCL28 and AtSMOS1 together at the same genomic sites.**

**(a)** Co-occurrence of ChIP-seq peak of SCL28 and AtSMOS1 at the genome-wide scale. SCL28 and AtSMOS1 tag density was compared in the ±5 kb region around the SCL28 peaks.

**(b)** Motifs enriched in SCL28 targets. HOMER motif search identified major SCL28-associated motifs.

**(c)** Motifs enriched in AtSMOS1 targets. HOMER motif search identifies major AtSMOS1-associated motif.

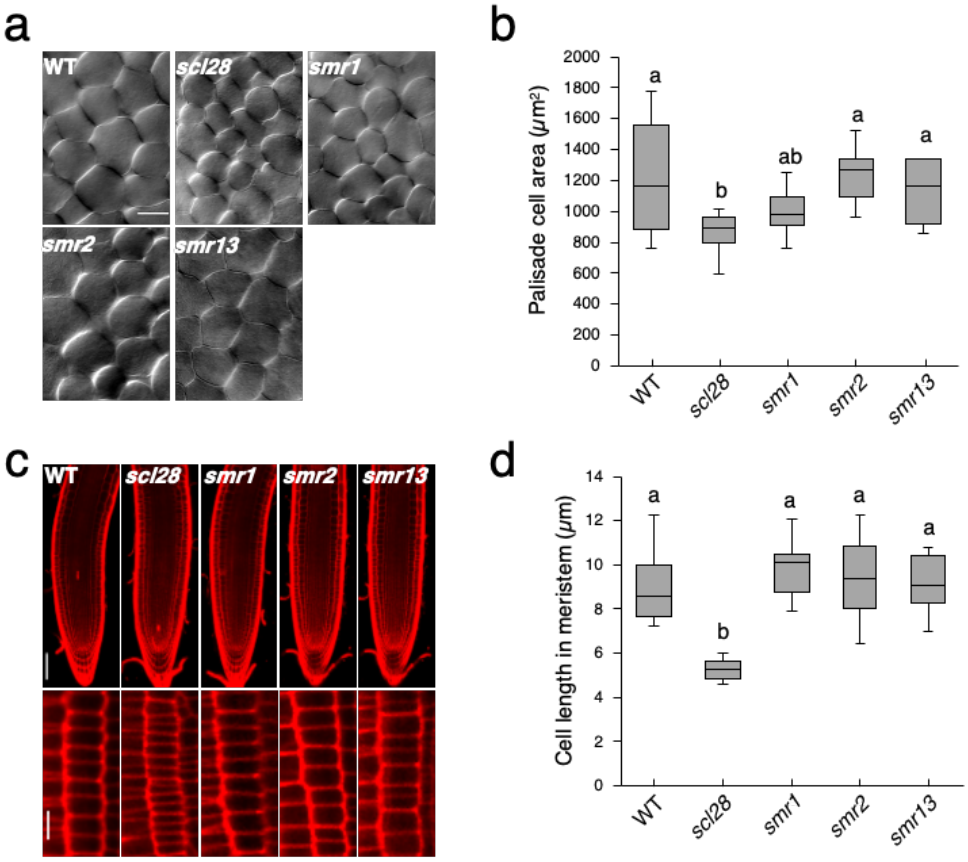


**Supplementary Figure 16**

**Cell size phenotype of *smr1*, *smr2*, and *smr13* plants.**

**(a)** First leaf pairs from plants at 22 DAS with indicated genotypes were cleared and observed with DIC microscopy. Scale bar indicates 30 $\mu$m.

**(b)** Quantification of palisade cell area in first leaf pairs from plants with indicated genotypes. Data are shown as boxplots (midline = median, box = IQR, whiskers = 1.5 × IQR). Different letters above boxplots indicate significant differences based on one-way ANOVA and Tukey's test, $P < 0.05$ ($n = 10$).

**(c)** Root meristems from plants at 7 DAS with indicated genotypes were stained by PI and observed with LSCM. Magnified views of cortical cell files are shown in lower panels. Scale bars indicate 50 $\mu$m (upper) and 15 $\mu$m (lower).

**(d)** Quantification of cell length in cortical cell files from plants with indicated genotypes. Data are shown as boxplots (midline = median, box = IQR, whiskers = 1.5 × IQR). Different letters above boxplots indicate significant differences based on one-way ANOVA and Tukey's test. $P < 0.05$ ($n = 10$).

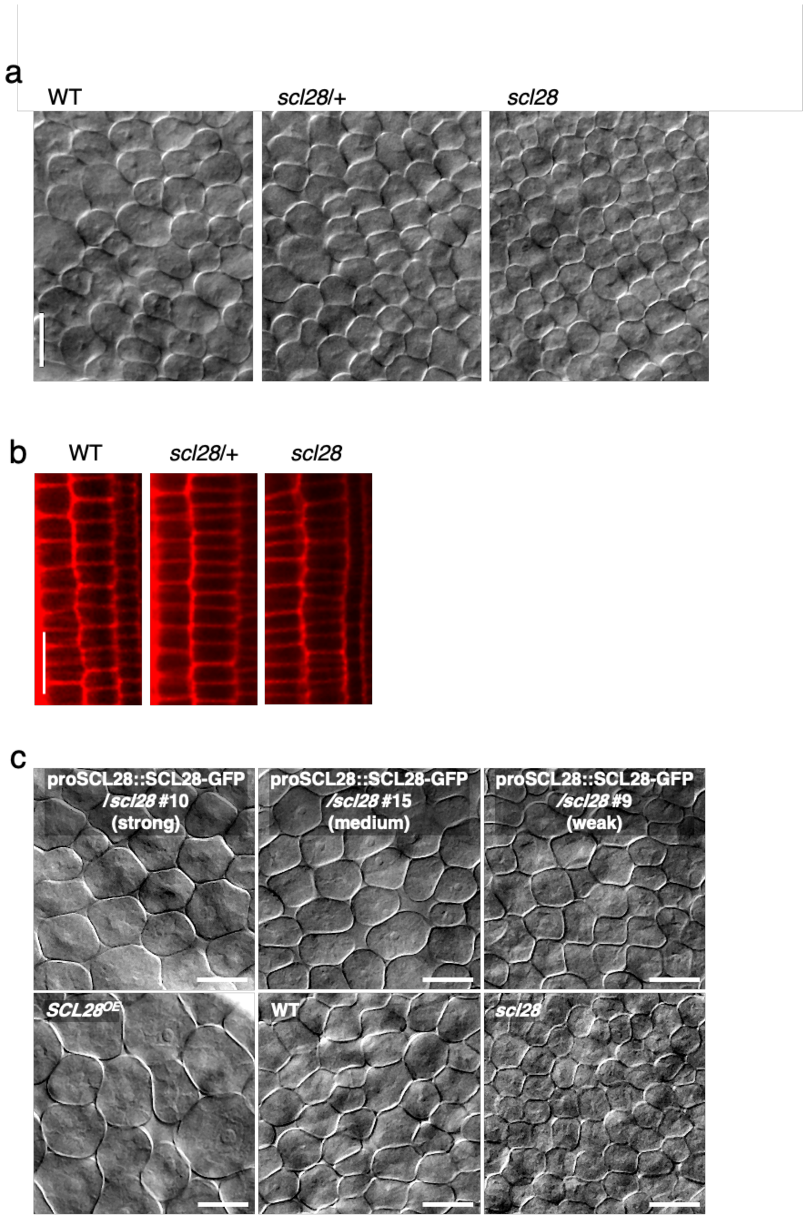


**Supplementary Figure 17**

**Cell size phenotype of plants heterozygous for *scl28* and those with moderately increased *SCL28* expression.**

**(a)** Images of palisade cells from WT plants and those heterozygous (*scl28*/+) or homozygous (*scl28*) for *scl28*. DIC microscopy observations were made for cleared first leaf pairs from plants at 10 DAS. Scale bar indicates 25 $\mu$m.

**(b)** Images of cortical cells in root meristems from WT, *scl28*/+, and *scl28* plants. Meristems of primary roots were stained by PI and observed with LSCM.

**(c)** Images of palisade cells from different transgenic lines of proSCL28::SCL28-GFP under *scl28* background with various expression levels (strong, medium, and weak). For comparison, WT, *scl28* and *SCL28*$^{OE}$ plants were also analyzed. DIC microscopy observations were made for cleared first leaf pairs from plants at 10 DAS with indicated genotypes.

### Supplementary Table 1

Primers used in this study

| Primer name | Purpose | Sequence |
|---|---|---|
| AtE1M gene-5/CACC | Construction of proSCL28::GUS and proSCL28::SCL28-GFP | CACCTTTGTCACGGCTTTAAACGCTTC |
| AtE1M gene-3 | Construction of proSCL28::SCL28-GFP | AATATAATGGGCCGACCTCATAC |
| AtE1Mp3-1 | Construction of proSCL28::GUS | AAGTCGACAACCCAATTCAAGAGATGGTCTAC |
| AtE1Mpro_dMSA5 | Site-directed mutagenesis of proSCL28::GUS | AAATTGGTGACCAATTGTGAATTGTCGGAGAAATATGACCAATGGGAG |
| AtE1Mpro_dMSA6 | Site-directed mutagenesis of proSCL28::GUS | AATTGGTCACCAATTTTCTACCAATTGATTTTAAAGATCCATTGGCTC |
| AtSMOS1 CDS-F1/CACC | Construction of pro35S::AtSMOS1 | CACCATGGCGTCGGTGTCGTCGTCGGAT |
| AtSMOS1 CDS-R1wStop | Construction of pro35S::AtSMOS1 | TCATTTCTCTTGTGGGAGGTA |
| SMR2p-Sma1-F | Construction of proSMR2::LUC | CACCCCCGGGAACTCCTTCGGCATCTTTGTTT |
| SMR2p-BamH1-R | Construction of proSMR2::LUC | CCCGGATCCGGTCACATGGATGTGAAAGTTT |
| AtE1M-5 | Construction of proRPS5A::SCL28-GFP | CACCATCTCTTGAATTGGGTTAGGTAG |
| GFPstop-R | Construction of proRPS5A::SCL28-GFP | TTACTTGTACAGCTCGTCCATGCCG |
| AtSMOS1 gene attB1_F | Construction of proAtSMOS1::AtSMOS1-GFP | GGGGACAAGTTTGTACAAAAAAGCAGGCTAGACTTATGCAATACTGTTGG |
| AtSMOS1 gene attB2_R | Construction of proAtSMOS1::AtSMOS1-GFP | GGGGACCACTTTGTACAAGAAAGCTGGGTTCTGGGTAATAGGATTCAGTT |
| SMOS1gene-EGFP_Cfu_F | Construction of proAtSMOS1::AtSMOS1-GFP | GACGAGCTGTACAAGTGAGCCGTTCCCTTTAGACTTTATG |
| SMOS1gene-EGFP_Cfu_R | Construction of proAtSMOS1::AtSMOS1-GFP | GCCCTTGCTCACCATTTTCTCTTGTGGGAGGTAGCTG |
| AtE1M_over insertion_qF | qRT-PCR for SCL28 | GCCTTCAAACAGAGGAATCTGG |
| AtE1M_over insertion_qR | qRT-PCR for SCL28 | TTGGTAAGCTTCATGGGAGCTC |
| CYCB1;2-qFW | qRT-PCR for CYCB1;2 | GAATATGGTTCACTTCCTTGC |
| CYCB1;2-qRV | qRT-PCR for CYCB1;2 | CTGCAATGTATCAGTCCAAGC |
| UBQ5_F | qRT-PCR for UBQ5 | CTTGAAGACGGCCGTACCCTC |
| UBQ5_R | qRT-PCR for UBQ5 | CGCTGAACCTTTCCAGATCCATCG |
| KNOLLE-Q4 | qRT-PCR for KNOLLE | TGATGGTTGAATCGCAAGGTGAAC |
| KNOLLE-Q3 | qRT-PCR for KNOLLE | TGCAGTCTTCAGCTCATTAGCTC |
| SIM_F1 | qRT-PCR for SIM | TCTTCCGACCACAAGATTCC |
| SIM_R1 | qRT-PCR for SIM | TCTTGCAAGATCTGATGCCG |
| SMR1_qF3 | qRT-PCR for SMR1 | CACCCACATCCCAAGAAC |
| SMR1_qR3 | qRT-PCR for SMR1 | GACGGAGGAGAAGAAACG |
| SMR2_F3 | qRT-PCR for SMR2 | CAAGATTGTCCAAAGATCTTCGG |
| SMR2_R1 | qRT-PCR for SMR2 | GGCACTATTACTCCTTCGTTTC |
| SMR3_qF | qRT-PCR for SMR3 | CGATCACAAGATTCCGGAGGTG |
| SMR3_qR | qRT-PCR for SMR3 | CGGCTCAGATCAATCGGTATGC |
| SMR4_qF2 | qRT-PCR for SMR4 | TGGTGGTGAGAAAACGAGATCC |
| SMR4_qR2 | qRT-PCR for SMR4 | AGGCTGTGCGTAGAACAAAG |
| SMR5_F | qRT-PCR for SMR5 | CGTGATGATTGCCGGATACC |
| SMR5_R | qRT-PCR for SMR5 | AAAATATCCCTTCTTCGGTGGTTC |
| SMR6_qF2 | qRT-PCR for SMR6 | TTTCGATTCGGGCTTCGTTG |
| SMR6_qR2 | qRT-PCR for SMR6 | TCTTCGTTTCCTTCGCTGTC |
| SMR7_F1 | qRT-PCR for SMR7 | CAGAGAATTAGACACCGATG |
| SMR7_R1 | qRT-PCR for SMR7 | CGTGGGAGTGATACAAATTC |
| SMR8_qF3 | qRT-PCR for SMR8 | AAACCGTCGTTGAAGTGCAG |
| SMR8_qR2 | qRT-PCR for SMR8 | GGGATCAGAGTCGTGAAAACAG |
| SMR9_qF2 | qRT-PCR for SMR9 | AAAAGGTGGCGCAAAACTGC |
| SMR9_qR2 | qRT-PCR for SMR9 | GTTGACCAAGTGCGAAAACG |
| SMR10_qF | qRT-PCR for SMR10 | GCAAAGAAGGAGCAACCGTCAAG |
| SMR10_qR | qRT-PCR for SMR10 | CGGTGGACAAATTCTTGGCATCG |
| SMR11_qF | qRT-PCR for SMR11 | CTGCTTCGATCTCGGATTGTGTT |
| SMR11_qR | qRT-PCR for SMR11 | GACGAAGGAGGCGGTGTTTTAC |
| SMR12_qF | qRT-PCR for SMR12 | GGTATGTCGGAGACGAGCTTGA |
| SMR12_qR | qRT-PCR for SMR12 | GAGTCGGTGTCTTGAACCCATCA |
| SMR13_qF | qRT-PCR for SMR13 | GAGTCTCCTGTAAAGATCCCAG |
| SMR13_R2 | qRT-PCR for SMR13 | TAGCTTTTGGCTTTCTCGGC |
| SMR14_qF1 | qRT-PCR for SMR14 | AACCAAACCGAGCCGAAAAG |
| SMR14_qR1 | qRT-PCR for SMR14 | GTGTTGATGTTGTTGTGTTGAGG |
| SMR15_qF | qRT-PCR for SMR15 | AATGCGTCATCACCGGAATC |
| SMR15_qR2 | qRT-PCR for SMR15 | TGGTGGCGAAAAGAACTCTC |
| SMR16_qF | qRT-PCR for SMR16 | GCCTTCAAACAGAGGAATCTGG |
| SMR16_qR | qRT-PCR for SMR16 | TTGGTAAGCTTCATGGGAGCTC |
| UBQ(-253/-32)_F | ChIP-qPCR for UBQ10 promoter | AATAAACGGCGTCAAAGTGG |
| UBQ(-253/-32)_R | ChIP-qPCR for UBQ10 promoter | ACGAGGACGACTAGGTCACG |
| pSMR2_(-567)_F | ChIP-qPCR for SMR2 promoter (distal) | GCGAAGGAGCGAATAATTCC |
| pSMR2_(-392)_R | ChIP-qPCR for SMR2 promoter (distal) | CGGAGAAGCTTATCCAAATTAATG |
| pSMR2_(-177)_F | ChIP-qPCR for SMR2 promoter (proximal) | GATTAATTCATACATGTACATTTTCGC |
| pSMR2_(-10)_R | ChIP-qPCR for SMR2 promoter (proximal) | GATGTGAAAGTTTCGTGGGC |
| SMR2_F3 | ChIP-qPCR for SMR2 CDS | CAAGATTGTCCAAAGATCTTCGG |
| SMR2_R1 | ChIP-qPCR for SMR2 CDS | GGCACTATTACTCCTTCGTTTC |
